# Amplified cortical neural responses as animals learn to use novel activity patterns

**DOI:** 10.1101/2022.07.10.499496

**Authors:** Bradley Akitake, Hannah M. Douglas, Paul K. LaFosse, Manuel Beiran, Ciana E. Deveau, Jonathan O’Rawe, Anna J. Li, Lauren N. Ryan, Samuel P. Duffy, Zhishang Zhou, Yanting Deng, Kanaka Rajan, Mark H. Histed

**Author notes:** denotes equal contribution (Bradley Akitake, Hannah M. Douglas).

## Abstract

Cerebral cortex supports representations of the world in patterns of neural activity, used by the brain to make decisions and guide behavior. Past work has found diverse, or limited, changes in the primary sensory cortex in response to learning, suggesting the key computations might occur in downstream regions. Alternatively, sensory cortical changes may be central to learning. We studied cortical learning by using controlled inputs we insert: we trained mice to recognize entirely novel, non-sensory patterns of cortical activity in the primary visual cortex (V1) created by optogenetic stimulation. As animals learned to use these novel patterns, we found their detection abilities improved by an order of magnitude or more. The behavioral change was accompanied by large increases in V1 neural responses to fixed optogenetic input. Neural response amplification to novel optogenetic inputs had little effect on existing visual sensory responses. A recurrent cortical model shows that this amplification can be achieved by a small mean shift in recurrent network synaptic strength. Amplification would seem to be desirable to improve decision-making in a detection task, and therefore these results suggest that adult recurrent cortical plasticity plays a significant role in improving behavioral performance during learning.

## Introduction

Sensorimotor decision-making involves patterns of neural activity which propagate through the neural circuits of many brain areas and are changed by those circuits. The sets of neural computations involved in sensory decision-making have not been fully determined^1–4^, but some principles have been identified. One basic neural computation is representation, storing information about the sensory world in patterns of activity, as is observed in many cerebral cortical areas. Another is decision, or readout, in which representations are transformed or categorized by circuits into forms suitable for action^5, 6^.

There is substantial evidence that sensory cortical representations can be modified by activity^7–11^, but it is less clear whether cortical response changes constitute the computational change that leads to improved behavior with learning. Studies in humans and animals have reported varied effects of learning on visual cortical responses, including increased activity after visual training^12–15^, selective suppression of activity^16^, decreased variability of visual selectivity response properties after training^17–19^, and activity changes that disappeared once early learning has ended^20^. Some learning studies have found improvement in primary sensory representations^19, 21–23^, along with changes in anticipatory and other signals^18, 24^. Other studies in primary visual cortex (area V1) have found little task-relevant change^16, 25^, but found changes in higher visual areas like V4^26, 27^. Thus, it has been unclear whether a major substrate of visual sensory learning is representational improvement in V1, such as increased gain or selectivity, or whether the principal changes are readout changes, perhaps in downstream areas.

One reason it has been difficult to delineate the neural computations underlying sensory decisions is that neurons and brain areas are highly interconnected, and sensory stimuli change activity in many brain areas^28–30^. Thus, changes in neural activity that are observed in one cortical area may be inherited from input regions, and indeed cognitive factors like attention or arousal can modulate visual activity before it arrives at the cortex^31^. One way to isolate cortical representations from downstream readout computations is to use stimulation-based behavioral paradigms. Using electrical or optogenetic stimulation methods, entirely novel (non-sensory, or ‘off-manifold’)^32, 33^, activity patterns can be introduced in a chosen brain region. Using such novel patterns is a way to explore the limits of cortical plasticity, as they are dissimilar from normal sensory patterns.

Here, to isolate representational changes that occur as animals improve on a task, we study V1 neural changes as mice learn to use a new cortical representation induced with optogenetic stimulation. Animals show dramatic improvements in behavior as they learn, with detection thresholds improving at times over several orders of magnitude during weeks or months of learning. Alongside the behavioral improvements, cortical neurons produce larger responses to the same optogenetic input. Thus, learning enables a fixed input to produce an increasingly large response in the V1 network, presumably by some adjustment of local, recurrent circuitry^34–36^. The results imply that this learning leads to local changes in representations by increasing recurrent amplification in V1.

## Results

We trained animals to detect neural activity evoked by optogenetic stimulation, and measured cortical responses during learning with 2-photon imaging. We implanted a 3 mm optical glass window over V1 and used multiple viral injections in layer II/III to express an opsin (soma-targeted ChrimsonR; stChrimsonR; excitatory neurons, AAV9-FLIP/DIO in Emx1-Cre mouse line)^37–39^, and for 2-photon imaging, a calcium indicator (jGCaMP7s or 8s; all neurons; AAV9-hSyn)^40–42^.

We delivered optogenetic stimulation light through the objective (combined into the light path via a dichroic; Methods; Figure 1A) which robustly activates stChrimsonR-expressing neurons throughout layer II/III (∼500 µm diameter light spot at cortical surface; Figure S5 and Methods;^43^).

**Figure 1.**
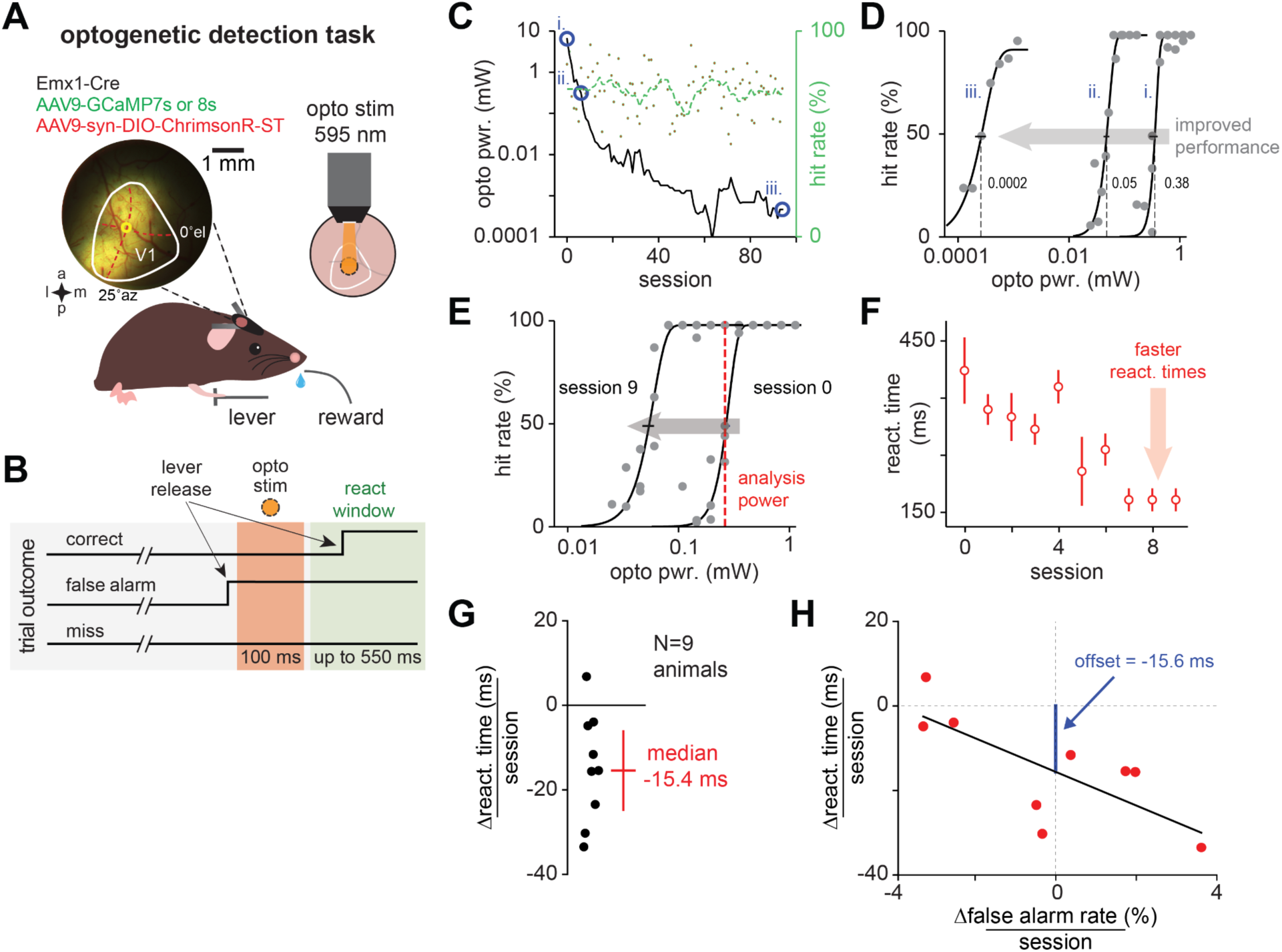
Mice gradually learn to report direct optogenetic stimulation of V1 excitatory neurons. (**A, B**) Task schematic. Animals release a lever when they detect the optogenetic stimulus (opsin: soma-targeted stChrimsonR in excitatory neurons). Only rapid lever releases (between 50-550 ms post-stim) were scored as correct (release before this window: false alarm, releases later: miss). (**C**) Example of long-term optogenetic learning. Blue circles, i-ii (7 sessions): initial fast drop in stimulation power required to hold performance constant (green: hit rate roughly constant, >70%; Results), ii-iii: longer phase of behavioral improvement (80 sessions). (**D**) Psychometric curves showing stimulation power decrease (curves from days shown by i, ii, and iii in 1**C**. Small gray text, threshold power in mW. Leftward shift signifies improved performance: gray arrow). Power threshold of the final session was three orders of magnitude lower than threshold of first session (0.38, 0.0002 mW: i, iii). (**E**) Psychometric curves covering the initial phase of optogenetic learning (same animal from **C,D**, sessions 0 and 9). Red dotted line: common power across sessions used for reaction time analysis. (**F, G**) Reaction times in response to optogenetic stimulation get shorter with learning (first 10 sessions of optogenetic learning, **F**: N = 1 animal, errorbars: SEM over trials, **G**: N = 9 animals, each point: regression slope for one animal, power shown by red line in **E**, Methods; errorbar: IQR = 18.6, p < 0.01). (**H**) Change in reaction time cannot merely be explained by change in false alarm rate, a proxy for response criterion^44^ (black line: linear regression, slope −3.98, p = 0.002, blue line: negative change in reaction time even at zero false alarm rate change, offset −15.6, p = 0.02.) Here and below, all errorbars: SEM unless otherwise specified.

Optogenetic detection training (N = 16 animals) occurred in two phases (Figure S1A,B). First, we trained animals to perform a sensory detection task. This was so they first learned the task demands (waiting for stimulus, lever press, etc.), reducing behavioral changes due to those effects as optogenetic learning progressed. We trained animals to respond to a small visual stimulus (monocular Gabor; 14° FWHM) until they performed the task with a stable psychometric threshold for three sessions (e.g., for animals imaged during behavior: training time 15-29 days, 23.6 ± 6.2 days, mean ± SEM, N = 3 animals). Next, we added an optogenetic stimulus (Figure 1; Figure S1; 0.5 mW at 595 nm), delivered at the same time as the visual stimulus. Over the course of several sessions, we removed the visual stimulus gradually by manually reducing visual stimulus contrast^45^. This made it more difficult to perform the task using the visual stimulus, but kept performance at approximately the same level as animals began to rely on the optogenetic stimulus (Figure 1A,B; Figure S1A,B). When contrast of the visual stimulus was zero, animals relied entirely on the optogenetic stimulus (2.3 ± 0.9 days after first optogenetic stimulus, mean ± SEM, animals used for imaging, N = 3; “session 0”). We confirmed that animals responded only to the optogenetic-evoked neural activity by moving the optogenetic spot during behavior to non-training locations within V1, which resulted in no behavioral responses (Figure S2A,B).

How similar are optogenetic responses to visual sensory responses? The optogenetic stimuli we use produce a different pattern of responses across the neural population than visual inputs, which activate cells based on their receptive field properties. However, in the temporal domain our optogenetic stimulation is more similar to visual responses, as optogenetic stimulation with the parameters we use modulates firing rates (measured with electrophysiology in^46^), and does not dramatically synchronize firing. This is consistent with the cortex operating as a recurrent network with reasonable strong excitatory-inhibitory coupling. In such a network, cortical neurons can fire irregularly, due to large amounts of recurrent input that lead to highly fluctuating membrane potentials^47–49^. Inputs then modulate the firing rate^43, 50, 51^ of the neurons — whose individual spike times are determined by the network-driven membrane potential fluctuations^52^.

### Optogenetic learning in a detection task

We found that animals dramatically increase their ability to detect the optogenetic stimulus – that is, the activation of V1 neurons — with practice. We collected psychometric curves during training sessions to track changes in animals’ perceptual sensitivity to the optogenetic stimulus (Figure 1C). Over the course of long-term training (∼90 sessions), we found that with practice animals’ perceptual thresholds dropped dramatically (Figure 1C). That is, animals needed less-strong stimulation over time to achieve the same level of performance. The observed rate of threshold change could be roughly separated into two phases, a phase that occurred within the initial ∼10 sessions of training after acquisition of the optogenetic task (Figure 1C,D: i and ii) and a slower phase over many additional sessions (Figure 1C,D: ii and iii). Below, we focus on the first six days of this initial learning phase for our experiments examining neural activity changes. In this initial phase, the threshold changes were large (Figure 1E, Δthresh. pwr. = −0.28 mW: 0.35, 95% CI [0.31-0.37], to 0.058 [0.052 - 0.063]).

The threshold changes were accompanied by decreases in reaction times. We compared reaction times for fixed stimulation powers across days (Figure 1F,G, median = −15.4 ms, IQR = 18.6, p < 0.01, over a subset of animals, N = 9, with common stimulation powers). The reaction time changes could not be accounted for by changes in animals’ false alarm rates (Figure 1H). While reaction times did change with false alarm rates, as expected due to changes in underlying perceptual criterion, reaction time changes remained after regressing out false alarm rate (Figure 1H).

### Responses of V1 to optogenetic stimulation are amplified by learning

We next imaged neural responses to stimulation during the process of learning. We measured neural responses in layer II/III during the first six optogenetic learning sessions, where learning is rapid (Figure 1C-F; Figure 2A). During this period, animals’ showed a greater than 50% drop in their optogenetic detection thresholds (Figure 2A, Δthresh. pwr. from session 0 to 5, −62 ± 10%, N = 4 animals, different cohort than in Figure 1).

**Figure 2.**
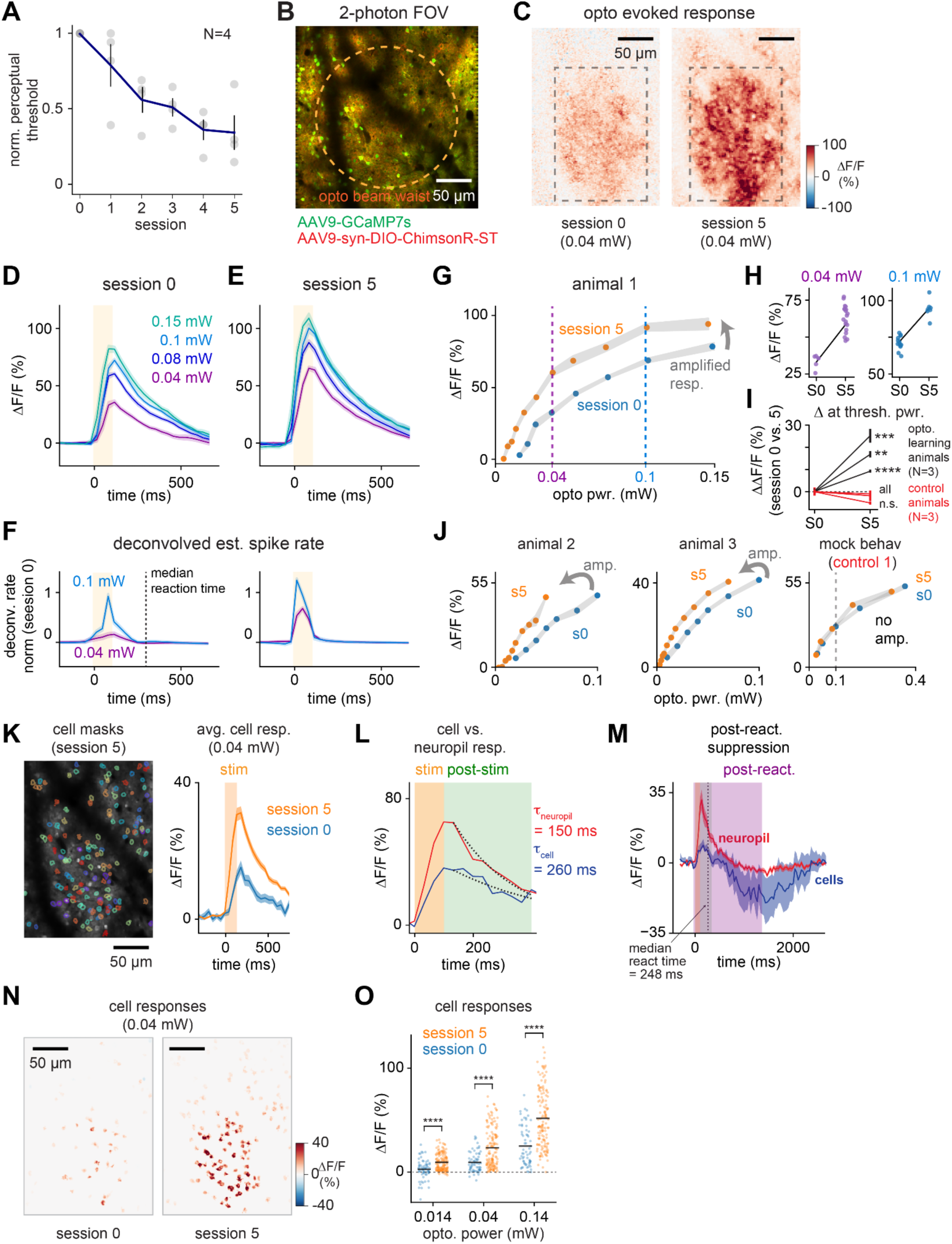
V1 responses to optogenetic stimulation are amplified by learning. (**A**) Animals improve optogenetic detection ability with practice (y-axis: threshold, stimulation power required for fixed detection performance; normalized to session 0, N = 4 animals imaged during learning, different cohort than Figure 1). (**B**) stChrimsonR, GCaMP7s expressed in layer II/III neurons (animal 1, shown over days in Figure S3). Orange circle: approx. stimulation beam waist (∼200 µm, Figure S5). (**C**) Neural response amplification after optogenetic learning (mean ΔF/F, 0.04 mW stimulation power, near psychometric threshold, animal 1, analysis from same animal for panels **D-H**). Grey box: region of interest (ROI) used for trial-by-trial ΔF/F analysis. (**D, E**) ΔF/F time courses before and after learning, matched stimulation powers. (**F**) Deconvolved signal (spike rate proxy; OASIS^53, 54^) shows spiking changes occur during stimulation (decay in **D-E** due to calcium dynamics, not spiking). (**G**) Average ΔF/F response across power levels (ROI shown in C). (**H**) Trial ΔF/F responses before and after learning (session 0 and 5: S0 and S5, ΔF/F in ROI, **C**; left: power near detection threshold, right: above threshold; each point one trial). (**I**) Normalized response change, all animals, with learning or control (change in ΔF/F, mean over trials ± SEM, at threshold power, ** p < 10^-2^, *** p < 10^-3^, **** p < 10^-4^, Mann-Whitney U test). (**J**) Same as **G**, for two additional animals plus an example control animal. (**K**) Left: cell masks (animal 2, session 5; found with suite2p^55^). Right: Mean cell responses before and after optogenetic learning. Orange box: optogenetic stim period (100 ms). (**L**) Example cell stimulation response (0.14 mW; more timecourses in Figure S6). Dotted lines: single-exponential fits to fluorescence decay (100-350 ms, green box). (**M**) Mean stimulation response in cells and neuropil is positive (left), but suppression is seen after animals’ responses (lever releases: dashed black line), purple: post-reaction averaging window. (**N**) Neuron responses to stimulation (during stimulation period: yellow in **L**,**M)** before and after learning (animal 2, near-threshold power for session 5, 0.04 mW). (**O**) Cell responses show widespread amplification with learning (each point: one cell, **** p < 10^-4^, unpaired t-test, session 0: N = 64, session 5: N = 142). N = 1 example animal in panels K-O.

To examine response changes with high signal-to-noise, we first averaged fluorescence responses over a large region of interest (Figure 2C). Imaging (Figure 2B) during optogenetic detection behavior then revealed clear stimulus-evoked responses that were strongly amplified over the course of training (Figure 2C-J).

This amplification could not be explained by shifts in the imaging plane or by changes in virus expression over sessions (Figure S3). It also could not be explained by tissue growth under the window or other optical degradation, over time or as a result of stimulation, as the effect we measured was in the opposite direction: an increase in responses to stimulation. However, to verify that optical changes did not account for the effects, we measured the effects of stimulation within each preparation at the imaging plane while not imaging, and used it to adjust stimulation power, finding that the amplification effects remained with and without this adjustment (Figure S4). Finally, as another check to rule out effects of imaging properties or expression contributing to this effect, we stimulated in control animals using matched mock training sessions, with the same imaging, stimulation, reward, optical window, and injection parameters as during training (Figure 2I,J; see also Figure S9 for similar control experiments using even higher powers). We found no amplification in this closely matched control (Figure 2I,J), arguing that the amplification we saw was indeed an increase in neural responses as a function of learning.

In principle it could have been that amplification was seen at some power levels but not others. We examined optogenetic-evoked responses and found that after learning, responses were amplified at all optogenetic power levels (Figure 2G-J), with strong effects both near the psychometric threshold (where behavior is tightly bound to stimulus perception) and also at above-threshold optogenetic stimulation powers (where trials are perceptually easy and performance is not stimulus-limited), where animals perform well. Though the magnitude of these changes varied somewhat across animals, we measured individually significant amplification in all learning animals and not in controls (Figure 2I; Figure S8).

We then examined single-neuron responses in an example animal (Figure 2K). We found that during the stimulation period, nearly all individual neurons (Figure 2K,N,O) as well as the surrounding neuropil (Figure 2L and Figure S6) showed positive responses. Thus, averaging neurons into large ROIs (Figure 2C-J) captures the effects seen in single cells, the positive responses across many neurons. The cell responses were amplified with learning (Figure 2O), and the amplification was seen across multiple powers (Figure 2N,O, mean change in ΔF/F = 6.7, 14.0, 26.6%; at 0.014, 0.04, and 0.14 mW; 95% CI [3.0 - 10], [9.5 - 19.5], [17.2 - 36.0]%), also consistent with the data from the large-ROI population measurements (Figure 2B-J). We also examined whether neurons showed any signs of suppression after stimulation^45^. We did find evidence for suppression (Figure 2M). However, this suppression was not part of the behavioral response or decision, as it occurred only after the animal made its behavioral response (Figure 2M and Figure S7, average reaction time for optogenetic learning animals 225 ± 23 ms, mean ± SEM, N = 3). This suppression timecourse is consistent with electrophysiological measurements of V1 excitatory optogenetic responses^46^. Those measurements show an initial positive transient in almost all neurons, followed in some excitatory cells by a suppressed steady state, effects that can be explained by coupling within the cortical recurrent network. In any case, for our 100 ms optogenetic pulses, we found the neural responses during the stimulation period were nearly entirely positive (Figure 2D-F,M-O), and further, these responses increased with learning.

The changes we observed in neural activity were smaller than the improvements seen in perception. Animals’ perceptual detection performance improved, and thresholds decreased, by a factor of approximately 2.7x after 6 sessions (i.e., power threshold was 37 ± 11% of session 0 levels; Figure 2A). In contrast, ΔF/F over the course of 6 sessions, measured at threshold stimulation power showed a 1.7x increase in ΔF/F over the large ROIs: session 0, 25.6 ± 7.4%, mean ± SEM across animals, session 5, 42.9 ± 10.9%, Figure 2I; and a 2.1x increase in mean cell peak ΔF/F, Figure 2O, 9.3% to 23.3%. Several caveats apply: the readout mechanism presumably sums across large numbers of neurons and thus may not be limited by the change in cortical responses we measure, and opsin saturation at high power may lead to greater changes in power than activity. However, the fact that behavior changes by a larger factor than cortical responses could potentially indicate that there is an improvement in the readout mechanism, occurring along with the amplification changes we see.

### The largest neural response changes happened from one day to the next, not within-session

While we observed significant increases in ΔF/F responses across experimental days, we found no evidence of increases within-session. In fact, we found a small decrease in responses to stimulation over the course of each experimental day (Figure S8, average ΔF/F change over 100 trials: −1.2% ΔF/F, 95% CI [-0.9 to 1.6]% ΔF/F, coeff. less than zero at p < 10^-13^, via linear regression over trials within day, estimated across animals and sessions, N = 3; Methods). Thus, it appears that optogenetic learning-related changes do not happen within the behavioral day, i.e., from one trial to the next. Instead, these data support that the major changes to neural responses occur outside of training, and may be driven by consolidation: changes in the brain in the hours between the experimental sessions.

### No amplification occurs with stimulation outside of the behavioral learning context

To determine if cortical amplification is dependent on learning, or might arise from repeated optogenetic stimulus alone, we performed a stimulation control in a mock behavioral context, and found no amplification (Figure 2I,J). That experiment was conducted with stimulation powers matched to those used during optogenetic learning (up to 0.5 mW, N = 3 animals). To determine if we could drive changes using stronger optogenetic stimulation, we increased stimulation power levels up to twice that used for behavior. We provided repeated optogenetic stimulation using a range of powers up to 1 mW (100 ms stimulation with ∼6 s interpulse interval, 1200 and 1500 repetitions, N = 2 animals, thus N = 5 total non-behaving controls). Even with higher stimulation powers we observed no changes in the optogenetic sensitivity of cells in the stimulated regions (Figure S9A,B). This result shows that amplification in response to these novel non-sensory stimuli requires an associative (behavioral) context.

### Statistics of visual responses are unchanged after optogenetic learning at both the training and control sites

Previous studies suggest that learning in visual perceptual tasks can lead to changes in the tuning properties of responsive neurons in mouse V1^19, 24^. However, it remains unresolved if these perceptual learning changes arise from plasticity in the local cortical networks, or if changes may be inherited from thalamic input pathways that could in principle adjust input strength, state, or synchrony^56–59^ to change cortical responses. Since optogenetic stimulation bypasses feedforward input from the thalamus, we asked whether the visual response properties of V1 neurons would change with optogenetic learning.

We imaged V1 neurons as mice were shown a series of visual stimuli before and after optogenetic learning (Figure 3A-D; Methods). We collected the responses of neurons at both the optogenetic training location (a V1 imaging site to which the visual stimulus was retinotopically matched), and an adjacent control location in V1 where stimuli were not delivered for optogenetic learning.

**Figure 3.**
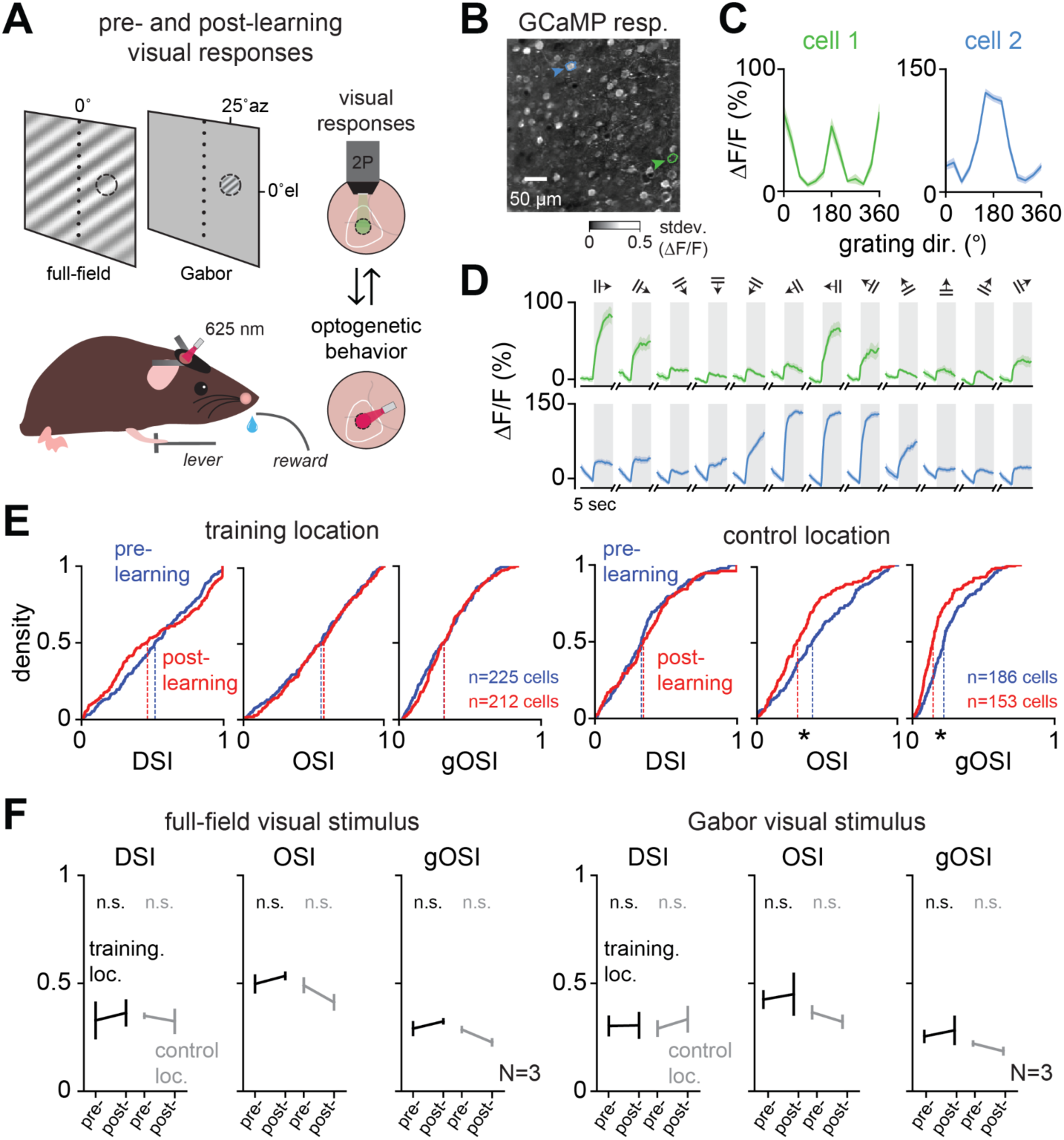
Visual response properties are unchanged after optogenetic learning. (**A**) Schematic of experiment (N = 3 animals, different animals than shown in Figure 2). Visual responses were measured before and after optogenetic learning (12-direction full-field drifting gratings or monocular Gabors, FWHM 12°) (**B**) Example pixel-by-pixel responses (gray: std. dev of ΔF/F over imaging frames.) (**C**) Tuning of two example cells (blue, green outlines in **B**). (**D**) Responses to visual stimulation across the 12 drifting grating directions, same cells in **C**. (**E**) Example (N = 1 animal) distributions of unitless indices for direction selectivity (DSI), orientation selectivity (OSI), and global orientation selectivity (gOSI; Methods), full-field stimulus; * p< 0.05, Kolmogorov-Smirnoff 2-sample test: p-values: training location, DSI: 0.06, OSI: 0.90, gOSI: 0.21; control location, DSI: 0.27, OSI: 7.8 x 10^-4^, gOSI: 5.7 x 10^-6^). (**F**) Summary of all visual response indices, both visual stimuli, pre- and post-optogenetic learning (mean ± SEM; n.s.: p > 0.05, N = 3 animals, unpaired t-test, pre-versus post-learning, also Figure S10).

Though we found some changes in visual tuning indices (Figure 3E) at both the optogenetic training and control locations before and after learning, these changes were inconsistent across animals and comparable in size between the training and control locations (Figure S10A). Across the population of animals, we found no significant mean changes in the visual response metrics (Figure 3F), nor in the magnitude of neural responsivity to visual stimuli (Figure S10B). The per-animal changes might perhaps arise from representational drift over time^60, 61^, potentially explaining why there was little mean change. The lack of mean change is consistent with the idea that recurrent network changes boost optogenetic responses, while leaving unchanged other dimensions of network response, as some overlap of responses must occur: many neurons respond to visual input (Figure 3E, Figure S10A), and with this viral expression approach, a majority of excitatory neurons express stChrimsonR^46^. Thus, while optogenetic learning leads to amplification of optogenetic responses, underlying visual response distributions and the overall structure of existing sensory representations remain intact.

### A network model shows amplification can be achieved by adjusting a minority of recurrent synapses

To understand how recurrent synapses might change to support the amplification we observed, we trained a recurrent neural network (RNN; Figure 4A) to show amplification. We trained the network in two steps, first to produce a response that mirrored an optogenetic input delivered to a fraction of cells (30%; matching previous expression data^46^, and Figures S4 and S5), and then to produce a response that was twice the size (Figure 4B). We only allowed changes in the recurrent connections, but not in the input and output weights. During training to produce amplification, many synaptic weights were adjusted, with a small positive shift in the population mean weight (Figure 4C,D, Figure S11, mean 5.8% ± s.d. 88% change). The stimulated neurons tended to strengthen their synapses onto other neurons (mean change 31.8%), while neurons that did not receive optogenetic input showed a small negative synaptic change (mean change −5.4%). The amplification in this recurrent model shows that synaptic strength changes, even when restricted to the local recurrent connectivity, can in principle support the amplification we observed.

**Figure 4.**
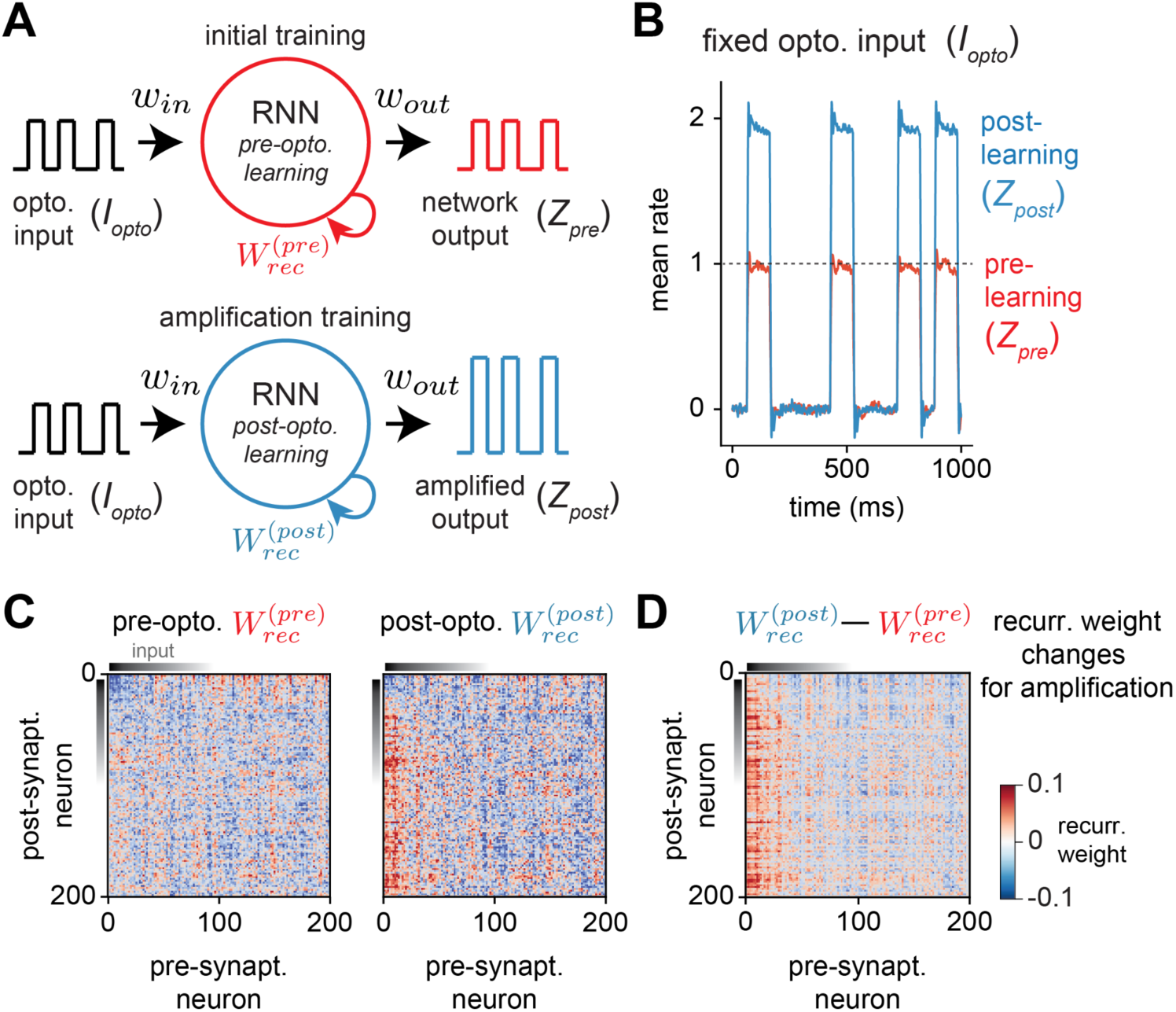
Network amplification for fixed optogenetic input arising from recurrent weight changes. (**A**) Two-step training of a rate-based RNN of 200 neurons with all-to-all connectivity (Gaussian distributed variance, g0 = 0.8; Figure S11; Methods). Fixed input (Win) and output (Wout) weights with 30% of neurons receiving optogenetic input (Iopto), a 100 ms pulse train with a variable rest interval up to 400 ms. Only recurrent weights were trained (W_rec_^(pre)^ and W_rec_^(post)^). Initial training (red): target output profile was Z_pre_ = I_opto_. Amplification training (blue): profile was Z_post_ = 2 * I_opto_, a fixed gain of 2. (**B**) Profiles of target optogenetic output mimics pre- and post-learning amplification (Z_pre_ and Z_post_, respectively). (**C**) Resultant weight matrices for initial training (W_rec_^(pre)^), and amplification training (W_rec_^(post)^). (**D**) Difference weight matrix (W_rec_^(post)^ – W_rec_^(pre)^) showing that amplification resulted in primarily positive weight changes across neurons receiving optogenetic stimulation.

## Discussion

In this work we examine the capacity of adult mouse primary visual cortex (V1) to undergo plastic changes in response to novel optogenetic stimuli over a few days of learning. We found clear evidence that neural responses to novel stimuli — optogenetic inputs applied directly to many cells — are amplified in V1, but only if those stimuli are made behaviorally-relevant. The changes in neurons’ responses over learning sessions mirrored the animals’ perceptual improvements. Responses to visual stimuli, which were not relevant for learning, did not show systematic changes, suggesting that the layer II/III cortical network was able to selectively amplify the input pattern created by optogenetic stimulation. Taken together, our results provide evidence for substantial plastic changes specifically in the primary visual cortex of the adult mouse brain that are linked to perceptual learning of a completely novel stimulus.

### Amplification is a desirable representational change for a perceptual detection task

In an optogenetic detection task, the principal neural computation that must be performed is a comparison between the activity evoked by optogenetic stimulation and spontaneous, ongoing activity. Therefore, the amplification of the optogenetic signal we found, an increasingly large spiking response to fixed input, seems to be the optimal way (assuming no major changes in the noise or variability in the population^62^) for the V1 recurrent network to adjust to improve task performance.

Other studies have found evidence for learning-related changes with optogenetic-stimulation tasks. Using a discrimination task and stimulating neurons in the somatosensory cortex (S1) with widefield (1-photon) optogenetics, Pancholi et al.^63^ found no evidence for amplification but did see other changes, including increases in response sparsity. Another study in S1 that used 1-photon stimulation learning^45^ found behavioral improvement, but did not examine neural changes during that learning. In the visual cortex, Marshel et al.^22^ trained animals to report activation of specific neural ensembles activated with 2-photon holographic stimulation. They found evidence for amplification in two different subnetworks (defined by intrinsic visual responses), but less-consistent changes for random-ensemble stimulation. In contrast, our work uses stronger widefield (1-photon) stimulation, and shows robust behavioral changes after learning that are accompanied by unambiguous V1 neural amplification.

The different effects seen in Pancholi et al. might be due to structural differences between V1 and S1 cortical circuits, or may be related to differences in task-specific computations. Their subjects were asked to discriminate between total stimulation intensity (low versus high number of optogenetic pulses), rather than discriminate or detect a specific pattern of activity.

Prior studies also disagree on interpretation, seemingly due to these differences in measurement of neural responses. For example, Dalgleish et al.^45^ hypothesize that the main neural changes relevant for behavior are happening downstream, outside the cortical area they stimulate (S1). Our work shows that there are clear changes occurring in V1 that support this optogenetic learning, and that those changes appear to be the optimal change to improve task performance.

### Readout changes and representational changes

Our results appear to help resolve a contradiction in recent optogenetic stimulation studies. Some studies have found animals can detect the activation of approximately 40 neurons, in somatosensory cortex (S1)^45^, and the olfactory bulb^64^. However, other work has found that only a subset of animals reported activation of similarly-sized groups of randomly selected V1 neurons^22^. While a possible explanation may be differences between brain areas, our data suggest a different explanation: that detection of randomly-selected small ensembles of neurons requires initial learning with stronger stimulation. The S1 and olfactory bulb studies initially trained animals using 1-photon (widefield) optogenetics, as we use here. Thus, these optogenetic results, along with electrical stimulation studies^65–70^ imply that, in many brain areas, animals can use completely novel, randomly-chosen patterns of neural stimulation, but to do so, learning must first be induced by strong stimulation of hundreds of neurons or more.

While we found significant changes in cortical representations during learning, it is possible that the readout mechanism improves as well. Our data might suggest there are changes in readout, beyond V1 changes in amplification, as we found larger improvements in behavioral performance than in cortical responses (percent changes in stimulation power needed to do the task vs. percent changes in neural responses; Figures 1 and 2), though interpretation is difficult due to potential opsin saturation and potential nonlinear or variability-dependent readout^62, 71^. Dalgleish et al. also provide evidence that readout changes occur in optogenetic-learning tasks: they found that high detection performance generalized across different stimulated patterns of cortical neurons. That is, after learning, animals did well at detecting the activation of not just a single trained subset of up to 100 neurons, but many different sets of up to 100 neurons. On the other hand, Marshel et al., who also stimulated randomly selected groups of up to approximately 100 neurons, found little generalization from one randomly-selected pattern to the next (their Figure 4I). Several differences might explain the divergent results: differences in cortical area, or difference in behavioral task: single-pattern detection vs. two-pattern discrimination. While our results show that cortical circuits can change with optogenetic learning, it is still possible that in some circumstances the decoding mechanism can also change during optogenetic learning.

The learning that we observed here seems likely to be a change in optogenetic sensitivity and not related to changes in movements. Our animals were pre-trained on a visual detection task before introducing the optogenetic stimulus (Figure S1A,B). Thus, the task demands and motor responses were fixed, and the only learning step needed was for animals to gain the ability to perceive and report the novel optogenetic activity induced in the cortex.

### Amplification happens via consolidation, with the largest changes outside sessions

Because we measured neural responses during task performance, we were able to determine whether amplification happened within the training sessions or developed from one day to the next. We found that within-session, there were small or negative changes in neural responses to a fixed stimulus (Figure S8), though there were consistent changes from one learning session to the next (Figure 2). While some decreases in response within-session could, in principle, be due to bleaching of opsin or indicator, the changes from one session to the next suggests that the major cortical network changes were happening outside sessions, perhaps as animals rested or slept. This is reminiscent of the consolidation that happens in motor learning, where a significant component of the motor improvement also appears to occur outside of the actual learning or practice repetitions^72, 73^.

Our physiological recordings found learning-related neural changes over the initial few days of optogenetic learning (5-6 days), consistent with previous reports^22, 45, 63^. However, we also measured continued improvement in optogenetic detection performance (without neural imaging) over many weeks to months of training (Figure 1). It seems possible that additional cortical amplification happens during this longer phase as well. This is supported by studies of long-term deafferentation, which have demonstrated that cortical responses can change over months or years to accommodate input changes^74, 75^.

### Pattern amplification in cortex due to recurrent connectivity

We found that optogenetic learning produced little change in the visual response properties of targeted neurons (Figure 3). In principle, the observed increase in cortical responses to the optogenetic stimulus could have arisen from changes outside the local cortical network that would not be due to modification of recurrent connections. These outside sources might be changes in top-down, higher-order thalamic (e.g., from the lateral posterior nucleus, LP / pulvinar) or neuromodulatory input that change the gain of V1 neurons. In addition, individual cells might change their intrinsic excitability^76^. However, were top-down input changes, intrinsic excitability, or neuromodulatory effects the dominant players, we might expect effects on visual responses as well. Theoretical work also shows that response amplification to a fixed input can be created in recurrent networks by adjusting the synaptic connectivity within the network^34, 35, 77^. Pattern completion observations in cortex^78^ are also consistent with response amplification, as amplification of a particular input pattern is closely related to completion, where a partial input pattern, via the recurrent network, induces larger responses in the neurons that compose the activity pattern. Finally, spinogenesis in motor cortex accompanies motor learning^79, 80^ and chronic optogenetic stimulation *in vitro* can also produce recurrent changes^81^. Together, along with the timecourse of the changes we saw, over the course of several days of practice, these observations suggest that changes in local recurrent cortical synapses are a likely mechanism for the learning-related neural changes we observed.

What circuit mechanism might gate, or enable, cortical recurrent plasticity, to allow changes during behavior but not for inputs presented outside a behavioral context? There is substantial evidence that inhibitory modulation is involved when such cortical network changes occur^9, 82–88^ and alternation of perineuronal networks, which surround many inhibitory neurons, participate in these synaptic changes^89–95^. Since the response changes we observed are dependent on animals performing a rewarded behavioral task, a compelling possibility is that task context or reward prediction signals trigger activation of inhibitory neurons, which opens the gate for plasticity, enabling changes to begin.

## Conclusion

How the cerebral cortex builds sensory representations for use in behavior is key to understanding brain function. Though the adult visual cortex is less plastic than the developing cortex^96–98^, our results – cortical amplification in response to completely novel artificial patterns of optogenetic input – provide key insights into how brains can adapt to behaviorally-relevant sensory information throughout our lifetimes.

## STAR★Methods

### Key resources table

| Reagent type (species) or resource | Designation | Source or reference | Identifiers | Additional information |
| --- | --- | --- | --- | --- |
| Genetic reagent (*M. musculus*) | Emx1-Cre | The Jackson Laboratory | RRID:IMSR_JAX:005628 | 21 total animals |
| Recombinant DNA reagent | AAV9-hSyn-FLEX-GCaMP6f | Addgene | #100833 |  |
| Recombinant DNA reagent | AAV9-hSyn-jGCaMP7s | Addgene | #104487 |  |
| Recombinant DNA reagent | AAV9-hSyn-jGCaMP8s | Addgene | #162377 |  |
| Recombinant DNA reagent | AAV1-hsyn-FLEX-ChrimsonR-tdTomato | Addgene | #62723 |  |
| Recombinant DNA reagent | AAV9-hSyn-DIO-stChRimsonR-mRuby2 | Addgene | #105448 |  |

## Resource availability

### Lead contact

Further information and requests for the resources should be directed to and will be fulfilled by the lead contact, Mark H. Histed.

### Materials availability

This study did not generate new unique reagents.

## Methods details

### Animals

All experimental procedures were approved by the NIH Institutional Animal Care and Use Committee (IACUC) and complied with Public Health Service policy on the humane care and use of laboratory animals. Emx1-Cre mice (Cre-recombinase targeted at the Emx1 locus^99^, Jax stock no. 005628, N = 21) were used for all experiments. N = 9 animals were used for optogenetic behavior without imaging (Figure 1), N = 4 for optogenetic behavior plus simultaneous 2-photon imaging (Figure 2), N = 3 for mock behavior with optogenetic stimulation only (Figure 2), N = 2 for non-behavior optogenetic stimulation (Figure S9), and N = 3 for visual stimulation before and after optogenetic behavior (Figure 3). Animals were housed on a reverse light/dark cycle.

### Cranial window implantation and viral injection

Mice were given intraperitoneal dexamethasone (3.2 mg/kg) and anesthetized with isoflurane (1–3% in 100% O_2_ at 1 L/min). Using aseptic technique, a titanium headpost was affixed using C & B Metabond (Parkell) and a 3 mm diameter craniotomy was made, centered over V1 (−3.1 mm ML, +1.5 mm AP from lambda).

Mice were injected with a pre-mixed combination of two adenovirus-mediated (AAV9) vectors for expression in the cortex, a functional calcium indicator (AAV9-hSyn-jGCaMP7s or - jGCaMP8s, viral titers 3.0 x 10^13^ and 4.1 x 10^13^ GC/ml respectively, final dilution 1:10) construct and a photoactivatable soma-targeted opsin construct (AAV9-hSyn-stChrimsonR-mRuby2, viral titer 3.2 x 10^13^ GC/ml, final dilution 1:8). Injections were made 150-250 µm below the surface of the brain for expression in layer II/III neurons. Multiple 300 nL injections were done at 150 nL/min to achieve widespread coverage across the 3 mm window. Animals were not reinjected.

A 3 mm optical window was then cemented into the craniotomy, providing chronic access to the visual cortex. Post-surgery, mice were given subcutaneous 72 hr slow-release buprenorphine (0.5 mg/kg) and recovered on a heating pad. Virus expression was monitored over the course of 3 weeks. We selected animals with good window clarity and high levels of virus co-expression (GCaMP and stChrimsonR) for behavior and imaging experiments.

### Retinotopic mapping

We determined the location of V1 in the cranial window prior to GCaMP or opsin expression using a hemodynamic intrinsic imaging protocol previously described in^100^. Briefly, we delivered small visual stimuli to head-fixed animals at different retinotopic positions and measured hemodynamic-related changes in absorption by measuring reflected 530 nm light. Imaging light was delivered with a 530 nm fiber-coupled LED (M350F2, Thorlabs). Images were collected through a green long-pass emission filter onto a Retiga R3 CCD camera (QImaging Inc., captured at 2 Hz with 4 × 4 binning). The hemodynamic response to each stimulus was calculated as the change in reflectance of the cortical surface between the baseline period and a response window starting 2–3 s after stimulus onset. We fit an average visual area map to the cortex based on the centroids of each stimulus’ V1 hemodynamic response.

These retinotopic maps were used during behavioral training to overlap the visual stimulus position in the right monocular hemifield with the imaging/optogenetic stimulation location in the V1. We found that the transition period between visual detection and optogenetic detection was facilitated by a strong overlap.

For measuring visual response properties, we further refined the visual position by measuring cellular responses in layer II/III with 2-photon imaging. Small oriented noise visual stimuli (14° FWHM) were presented at 9 locations (spaced by ±15° azimuth and ±10° elevation) in the right visual hemifield. The visual stimulus position that evoked the greatest response in the FOV was chosen for characterizing visual responses. We found that the strongest response was typically the center location, selected using the widefield hemodynamic map above.

### Behavioral task

Water-restricted mice (20-40 ml/kg/day) were head-fixed and trained first to hold a lever and release in response to a visual stimulus (Gabor patch; 14° FWHM, spatial frequency 0.1 cycle/degree), that increased contrast relative to a gray screen^100, 101^, and then to an optogenetic stimulus that directly activated layer II/III neurons in V1. Mice initiated behavioral trials by pressing and holding a lever for 400-4000 ms (according to a geometric distribution, to reduce variation in the stimulus appearance time hazard function, see^100^), and then the stimulus appeared for 100 ms in the animal’s right monocular hemifield. Animals had up to 550 ms to report the stimulus by releasing the lever. Because some minimum time is required to process the stimulus, we counted as false alarm trials those releases that occurred within 50-100 ms of the stimulus onset. Correct detection responses resulted in delivery of a 1-5 µL liquid reward (10 mM saccharine). We varied the liquid reward during training^101^, increasing reward after up to three consecutive correct trials, to decrease incentive for guessing^102^. Once proficient, reward volume did not fluctuate significantly across sessions.

All behavioral animals were first trained on a visual detection task (see task schematic, in Figure S1, and^100^). Once animals were performing well on the visual task and produced stable psychometric curves with low lapses for three consecutive sessions, we transitioned the animal to using the optogenetic stimulus by pairing each visual stimulus appearance with a fixed power (0.5 mW) optogenetic stimulation. During these transition sessions we lowered the contrast of the visual stimulus until animals could perform the task without the visual stimulus. The session where animals started behaving exclusively on the optogenetic stimulus was denoted session 0. During session 0 we generated the first psychometric curve for optogenetic stimulation. Analysis of data from session 0 came only from the part of trials where the animal was exclusively on the optogenetic stimulus. Subsequent behavioral sessions were started and conducted with only optogenetic stimuli. Animals used in behavior were not exposed to any other 1-photon stimulation outside of behavior and the craniotomy was kept covered by an opaque cap between sessions.

### Optogenetic stimulation

For optogenetic behavior experiments without simultaneous 2-photon imaging we delivered light through a fiber aimed at the cortical surface^100^. A fiber-coupled LED light source (M625F2, Thorlabs, peak wavelength 625 ± 15 nm, FWHM) was coupled via a fiber patch cable to a fiber optic cannula (400 µm core diameter, 0.39 NA, Thorlabs CFMLC14L02) cemented above V1. This method was used for long-term learning and control experiments with increased optogenetic stimulation outside of behavior (powers up to 1mW with 6.3 ± 1.7s between simulations, mean ± SD, N = 2).

For optogenetic behavior experiments conducted with simultaneous 2-photon imaging we activated stChrimsonR expressing neurons by passing 595 nm light (CoolLED pE4000 multispectral illuminator, 595 ±15 nm, FWHM) through the imaging objective to the surface of the brain. The illumination power was measured through the objective at the beginning of each session using a light meter (Newport 1918-C with a 918D-SL-OD3R detector) with a maximum of ∼0.5 mW.

### Analysis of behavioral data

Analyses were conducted in Matlab and Python. Optogenetic learning effects were characterized by analyzing data collected during animal behavior on the optogenetic stimulation detection task.

Reaction times were averaged across trials for each laser power group and for each training session. Linear fits were calculated for these data points across the start and end sessions in which each laser power group was present during the task. The slope of the linear fit indicated the change in reaction time per session for each laser power group. A mean change in reaction time per training session was then calculated across all laser powers for each animal. Changes in optogenetic detection sensitivity were analyzed by fitting cumulative Weibull functions to data from individual training sessions to estimate detection performance (hit rate) as a function of laser power. Quantifying thresholds with d’ (sensitivity) produces similar results to using hit rate in this task, as false alarm rates are nearly constant over time (false alarm hazard rate is near constant, see^100^). Threshold was the 50% point of the Weibull functions.

### 2-photon calcium imaging

2-photon calcium imaging was conducted using a custom microscope based on MIMMS (Modular In vivo Multiphoton Microscopy System, e.g.,^103^) components (Sutter Instruments, Novato, CA) with a Chameleon Discovery NX tunable femtosecond laser (Coherent, Inc.; Santa Clara, CA). Imaging was performed using a 16X water dipping objective (Nikon; Tokyo, Japan). A small volume of clear ultrasound gel (∼1 mL) was used to immerse the lens. Images of calcium responses (∼150-200 µm from the surface of the pia, layer II/III) were acquired at 30 Hz using ≤ 50 mW laser power for static imaging, and ≤ 15 mW for behavior at 920 nm.

### Analysis of imaging data

Raw 2-photon image stacks were downsized (512 rows to 256 rows) to facilitate handling of large datasets. For each behavioral session, frames were motion corrected using CaImAn^104^. Each imaging data set was baseline corrected to an estimated minimum pixel intensity, calculated as the minimum value in the average projection image across all frames from all trials prior to stimulus presentation (F_min_, a scalar). The minimum pixel intensity was subtracted from all pixels and all resulting negative values were set to 0.

For quantitative analyses we computed ΔF/F as (F-F_0_)/F_0_ at each pixel. F_0_ was taken over the 10 frames before each stimulus onset, and F_0_ did not systematically change over days (see also Figure S3). For statistical analyses F was taken as the frame 120 ms after the stimulus onset (frame 3 post-stimulation, near the peak response). For visual display of responses in entire frames, as in Figure 2C, F was taken over 0-270 ms after stimulus onset (frames 0-9 post-stimulation), and we computed ΔF/F as (F-F_0_)/F_div_, where F_div_ is F_0_ smoothed with a gaussian filter (sigma = 20 pix). Using a smoothed divisor image averages overall intensity in small regions of the image, yielding a form of local contrast adaptation. Image ROI fluorescent (F) activity traces were measured by calculating the average pixel intensity within a user-defined ROI, prior to computing ΔF/F for an ROI. Deconvolved calcium responses to estimate spiking activity for an ROI were calculated using the OASIS method with an autoregressive constant of 1^53^.

Segmented cell masks were identified using either Suite2p (for Figure 2)^55^ or CaImAn (for Figure 3)^104^ and their resulting calcium responses (F) were extracted. In order to quantify neuropil activity, we manually segregated cell bodies from their surrounding neuropil with non-overlapping masks (for Figure 2, details in Figure S6). We fit the fluorescence decays of cell bodies neuropil by a single exponential in a post-stimulation window (300 ms, starting 1 frame after cessation of optogenetic stimulation). Suppression effects were characterized in a 1.5 s post-reaction time window (starting 350 ms after optogenetic stimulus presentation, well after the median reaction time (∼250 ms) for the detection behavior).

Linear regression model for testing for effects of change between experimental days was OLS regression, using all trials on which the stimulus was successfully detected. Data was from N = 3 animals, N = 6 sessions for each animal, and 2633 total number of stimulation trials (all animals and sessions are shown in Figure S8, including the same analysis of N = 3 mock behavior control animals). Regression model equation: ΔF/F ∼ C(animal) * C(session) + stimulation_power_mw + trial_number + constant, where C(x) signifies a categorical or dummy variable. Full details of the model definition are in https://patsy.readthedocs.io/en/latest/.

We also tested for significant change in ΔF/F within-session by running the same model over each animals’ data, and found all three animals showed a negative change (trial number coefficient: −1.5, −1.1, −0.2% ΔF/F) though only two were significantly different from zero (p < 1 x 10^-12^, < 1 x 10^-6^, = 0.6, respectively).

Linear regression model for testing effects of optogenetic stimulation outside of behavior (results in Figure S9) was OLS regression from N = 2 animals, session 0 (S0) vs. session 6 (S6) via ANOVA. Regression model equation: ΔF/F ∼ C(power) + C(S0 v. S6), where C(x) indicates a categorical or dummy variable.

### Confirming optogenetic stimulation power between sessions

We measured the power of the stimulation LED light path immediately before each behavioral session. We also measured relative laser excitation power across days by measuring light collected by the PMTs during stimulation. The optogenetic blanking circuit operates the LED illuminator during the flyback phase of scanning image acquisition, and the refractory time of the blanking circuit leaves an up to ∼20 pixel artifact at the edges of the raw image stacks that scales with stimulation intensity. We used the mean pixel intensity change for this artifact to scale attenuated sessions and normalize stimulation powers across days (Figure S4), and our results were unchanged with and without this scaling, confirming we accurately measured stimulation power.

### Analysis of visual response properties

2-photon calcium imaging was performed directly before and after optogenetic learning to assess V1 neural responses at both training and control locations (an area with stable expression at least 200 µm away from training location). Visual stimuli were presented on a monitor positioned in front of the head-fixed animal at a 45° angle on the animal’s right side. The visual stimulus was either a full-field or Gabor patch (12° FWHM) drifting grating stimulus at 100% contrast presented in 12 different directions (30° increments). Stimuli were presented for 3 second durations (with 4 seconds between presentations) and were delivered in random order for a total of 25 repetitions of each stimulus direction. Gabor patch stimuli were displayed on the monitor at the visual field location corresponding to the retinotopic map at the training and control locations.

To assess potential changes in visual response selectivity, direction and orientation selectivity indices were calculated for each identified cell^105, 106^. First, tuning curves for each cell were calculated by averaging ΔF/F responses across the 3 second stimulus period across all repetitions for each of the 12 drifting grating directions. Direction selectivity indices (DSI) were measured as (R_pref_ - R_oppo_)/(R_pref_ + R_oppo_), where R_pref_ is the peak average response across the 12 directions and R_oppo_ is the average response at the opposite direction 180° away from the preferred direction. Orientation selectivity indices (OSI) were measured by first averaging responses from opposite pairs of directions (e.g., 0° and 180°, 45° and 225°) and calculating (R_pref_ - R_ortho_)/(R_pref_ + R_ortho_), where R_pref_ is the peak average response across the 6 orientations, and R_ortho_ is the average response of the orthogonal orientation 90° away from the preferred orientation. Last, a global OSI (gOSI) metric was calculated as 1 - CV (tuning curve) for each cell, where CV is the circular variance.

### Modeling

We trained a recurrent neural network (RNN) consisting of N = 200 units, whose input dynamics for the *i*-th neuron are given by:

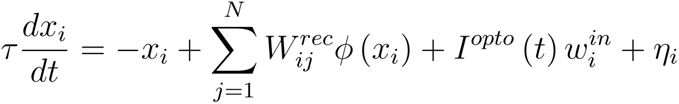

The readout of the network is defined as:

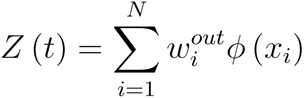

The transfer function of single units is *Φ(x)* = tanh*(x)*. The weights of the input pattern *w_in_* are positive and exponentially distributed for a fraction *p* = 0.3 of units, and zero otherwise. The readout weights are homogeneous and constant: 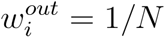. The initial recurrent weights 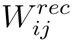, before any training, are independentlyis sampled from a random Gaussian distribution with mean zero and standard deviation 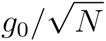^107^. The noise term *η_i_* is randomly sampled from a zero mean distribution with standard deviation 0.0005 at every time step.

We trained the recurrent weights 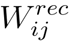 of the RNN using backpropagation-through-time (ADAM optimizer ^108^ in pytorch ^109^ such that the network readout matches a scaled version of the time-varying input *I^opto^ (t)*. The input and output weights remained fixed. In a first phase, mimicking the pre-learning response, we trained the network for 100 epochs such that *Z_pre_* = *I_opto_*, obtaining recurrent weights 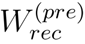. In a second phase, we trained the pre-learning network on 100 epochs to produce an amplified response, *Z_post_* = 2*I_opto_*, with recurrent weights 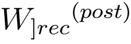. Parameters: τ = 10 ms, *g_0_* = 0.8, Euler integration timestep Δ*t* = 1 ms, learning rate 0.01.

To compute the normalized synaptic weight change in percent, we took the mean of the absolute value of weight across all synapses during the pre-training period, yielding a scalar value, and divided each synaptic weight by this scalar and multiplied by 100.

## Acknowledgments

We thank Victoria Scott for assistance with breeding and husbandry. A. Afraz, B. Averbeck, and members of the Histed laboratory for comments and discussion. This work was supported by the NIH Intramural program (ZIAMH002956) and NIH BRAIN Initiative (U19NS107464 and U01NS108683).

## Author contributions

B.A., H.D., P.K.L, L.R., and S.D. collected behavior and imaging data, with the help of Y.D. and A.L.. B.A., H.D., P.K.L., C.D., J.O., and M.H. performed data analysis. H.D., S.D., A.L., and Z.Z. prepared optical windows and did virus injections. M.B., B.A., K. R., and M.H. performed the modeling. B.A., H.D., P.K.L, and M.H. designed the experiments. B.A., H.D., P.K.L, J.O., M.B., and M.H. wrote the manuscript.

## Data availability

The datasets generated during the current study are available from the corresponding author on reasonable request. Data with plotting code are available at: https://github.com/histedlab/

## Supplemental Figures

**Figure S1.**
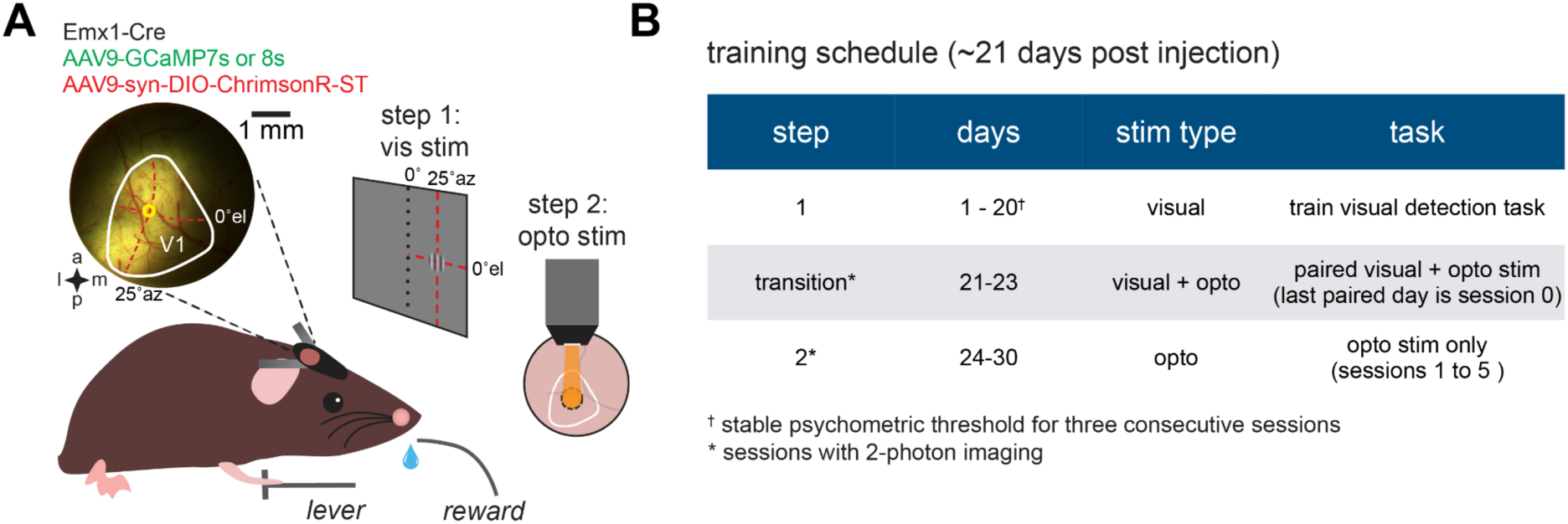
Training timeline for the optogenetic detection task, Related to Figures 1 and 2. (**A**) Schematic of 2- step protocol for behavioral training first on visual stimulus (step 1) then on optogenetic stimulation (step 2). The optogenetic stimulation location was aligned to the retinotopic location of the visual stimulation in V1. (**B**) Typical behavioral training schedule outlining the length of time for visual detection task proficiency and the steps to transition animals from the visual to the optogenetic stimulus (other statistics in Results). Visual detection proficiency was determined by animals achieving a stable psychometric threshold for three consecutive sessions (^†^). 2-photon imaging was conducted during the transition and step 2 sessions (*).

**Figure S2.**
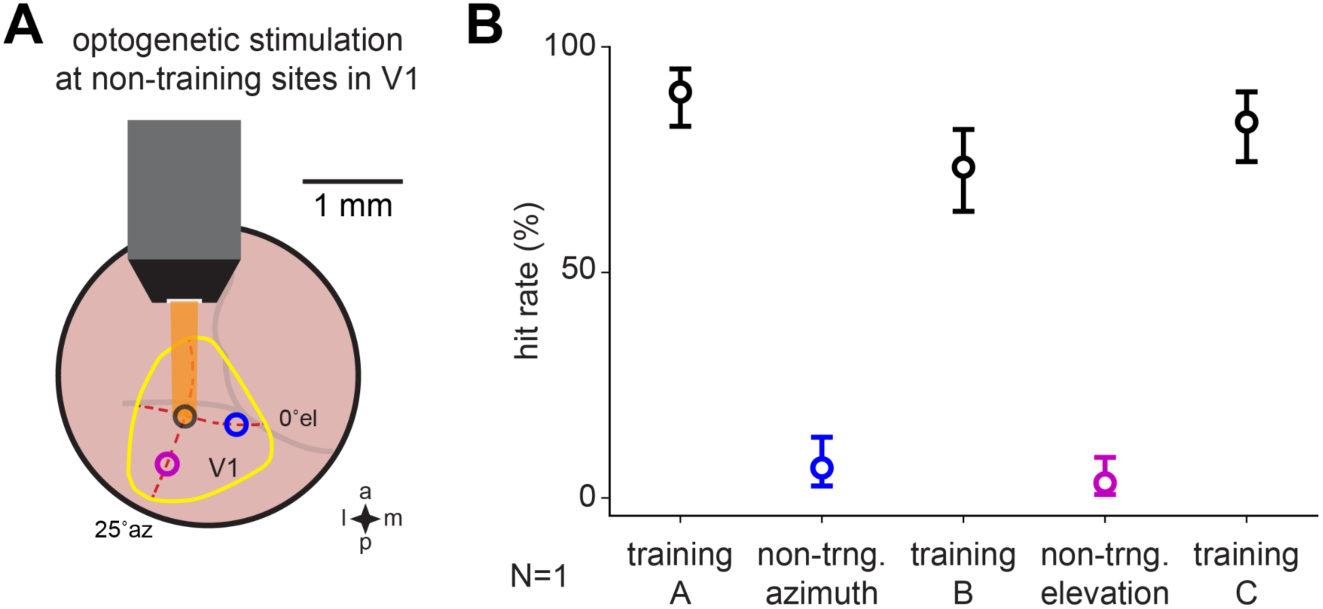
Animals detect and use the optogenetic-induced cortical activity; they do not detect stray light with their retinas, Related to Figure 1 and 2. (**A**) Schematic of experiment where we moved the stimulation light spot a small amount and found dramatic changes in behavior. This implies that animals’ behavior depends on cortical neural optogenetic activation. Black circle indicates optogenetic training location in V1 (yellow outline). After collecting a psychometric curve at the training location we moved the optogenetic stimulation ∼500 µm along the cortex, both in the visual-map-defined azimuth and elevation meridians (red dotted lines). At each of the shifted locations, blue and magenta circles, behavioral performance dropped and was recovered when we moved the stimulation back to the training location. (**B**) Detection hit rates in a trained animal during a session where the optogenetic stimulation location was sequentially moved for 30 trials each to and from non-trained locations in V1 (black, training A: 90.0 CI [82.4 - 95.1]%, training B: 73.3 [63.5 - 81.65]%, training C: 83.3 CI [74.5 - 90.1]%, blue, non-training azimuth change, 6.7 CI [2.7 - 13.4]%, magenta, non-training elevation change, 3.3 CI [0.7 - 9.0]%, hit rate ± Wald CI, N = 1).

**Figure S3.**
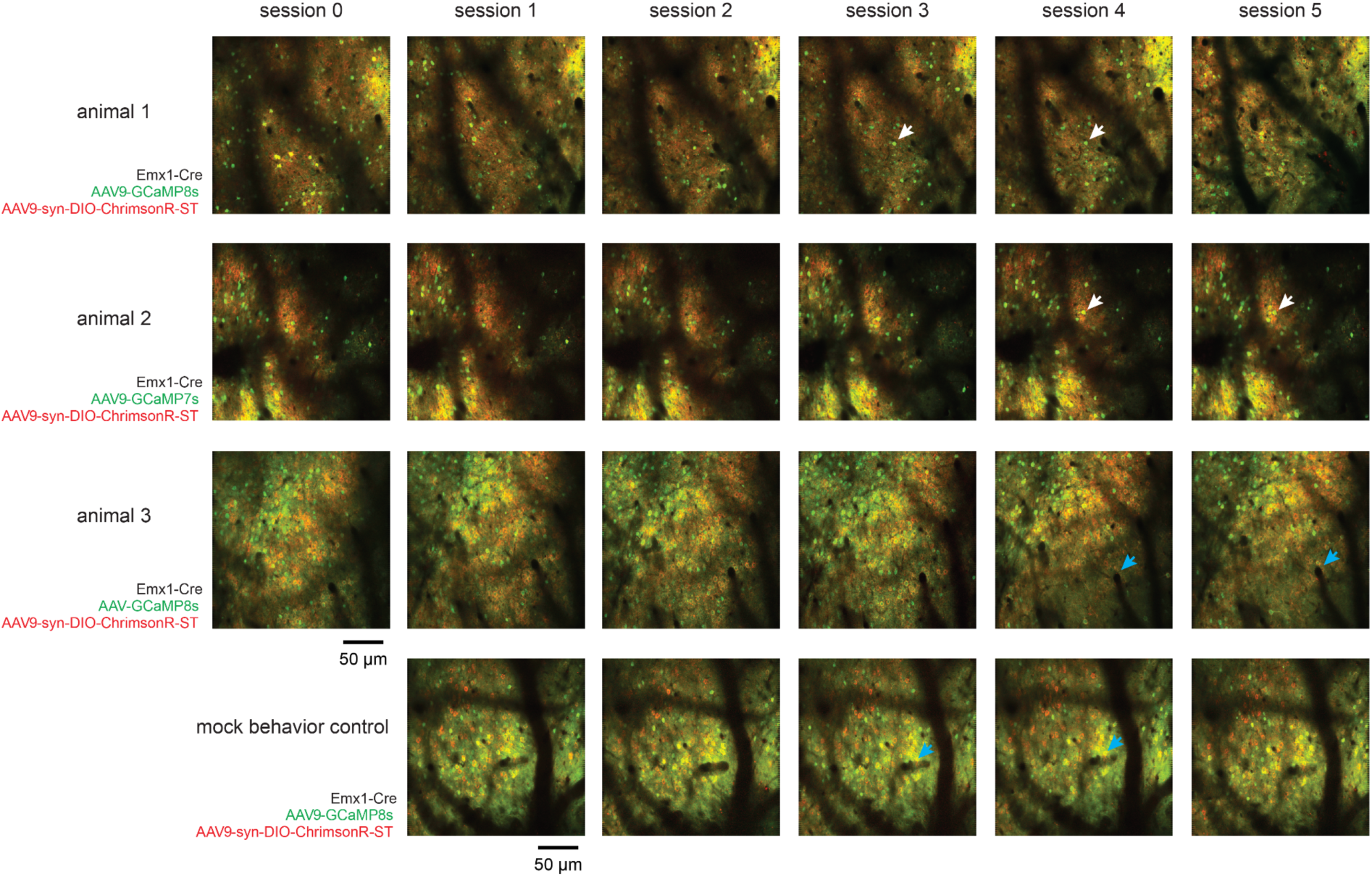
Imaging plane over sessions for optogenetic learning animals and mock behavioral control, Related to Figure 2. Genotypes and viral injections are listed for each animal tested. Imaging planes were aligned to reference GCaMP expressing cells (examples, white arrows) and vasculature patterns (examples, blue arrows) between sessions. All Red/Green images shown are 300 frame averages acquired with the same amplifier gain settings at 1000 nm excitation (∼35-45 mW). While some neurons differ from day to day, many of the same neurons were imaged across days.

**Figure S4.**
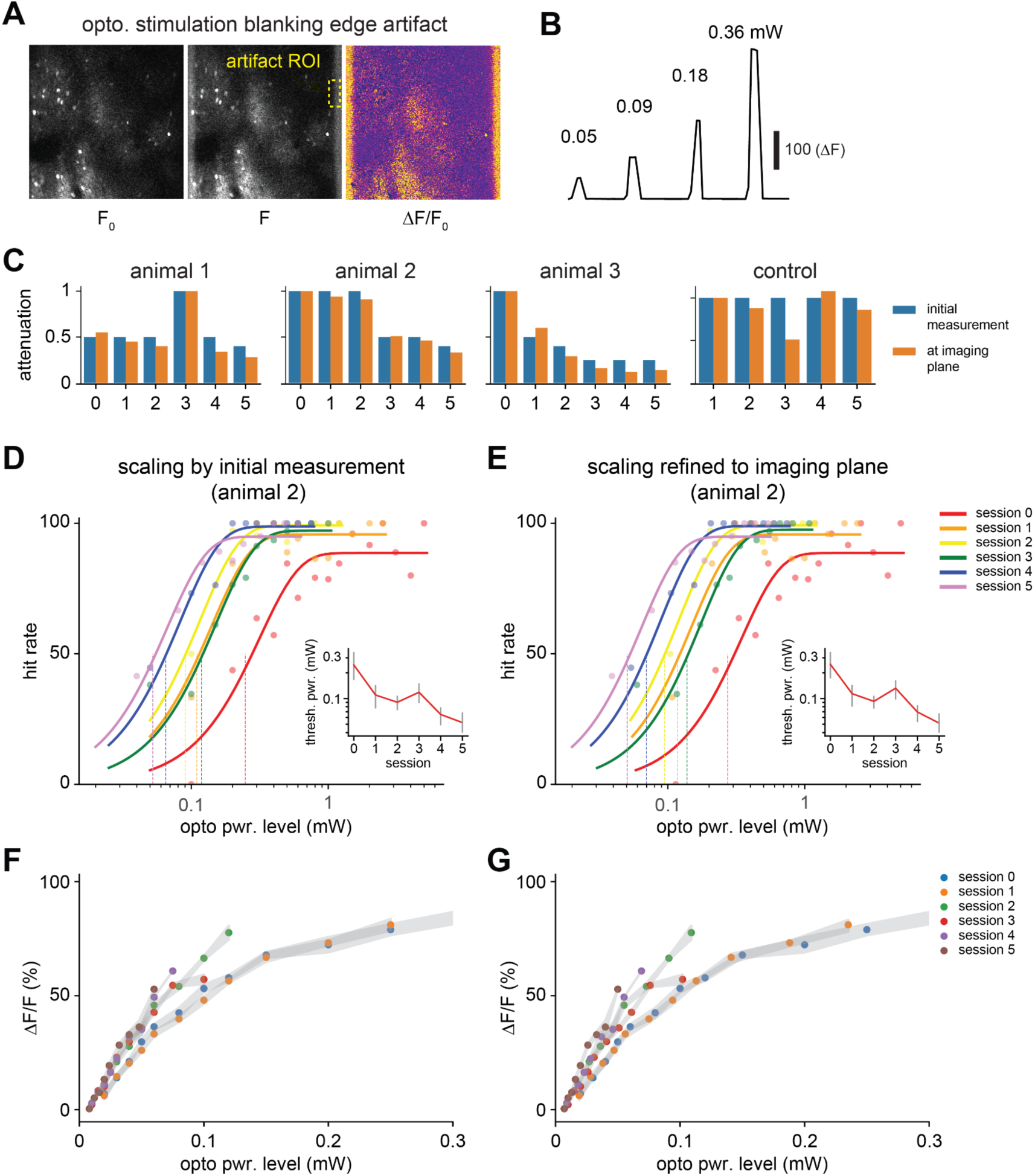
Optogenetic stimulation blanking artifact allows normalization of optogenetic power at the imaging plane between sessions, Related to Figure 2. (**A**) Optogenetic stimulation produces an ∼20 pixel edge artifact that is visible during imaging, as the optogenetic light source offset lasts a few microseconds into each imaging line after horizontal flyback. (**B**) Intensity of the edge artifact scales with applied optogenetic stimulation power. (**C**) Plots of attenuation based on initial measurement of power out of the objective and normalized scaling for all animals and control. (**D,E**) Normalized scaling refines the position of psychometric curves but does not change the order. Normalized scaling does not alter the relationship between threshold powers (insets). (**F,G**) Normalized scaling does not alter the relationship between ΔF/F and power over sessions.

**Figure S5.**
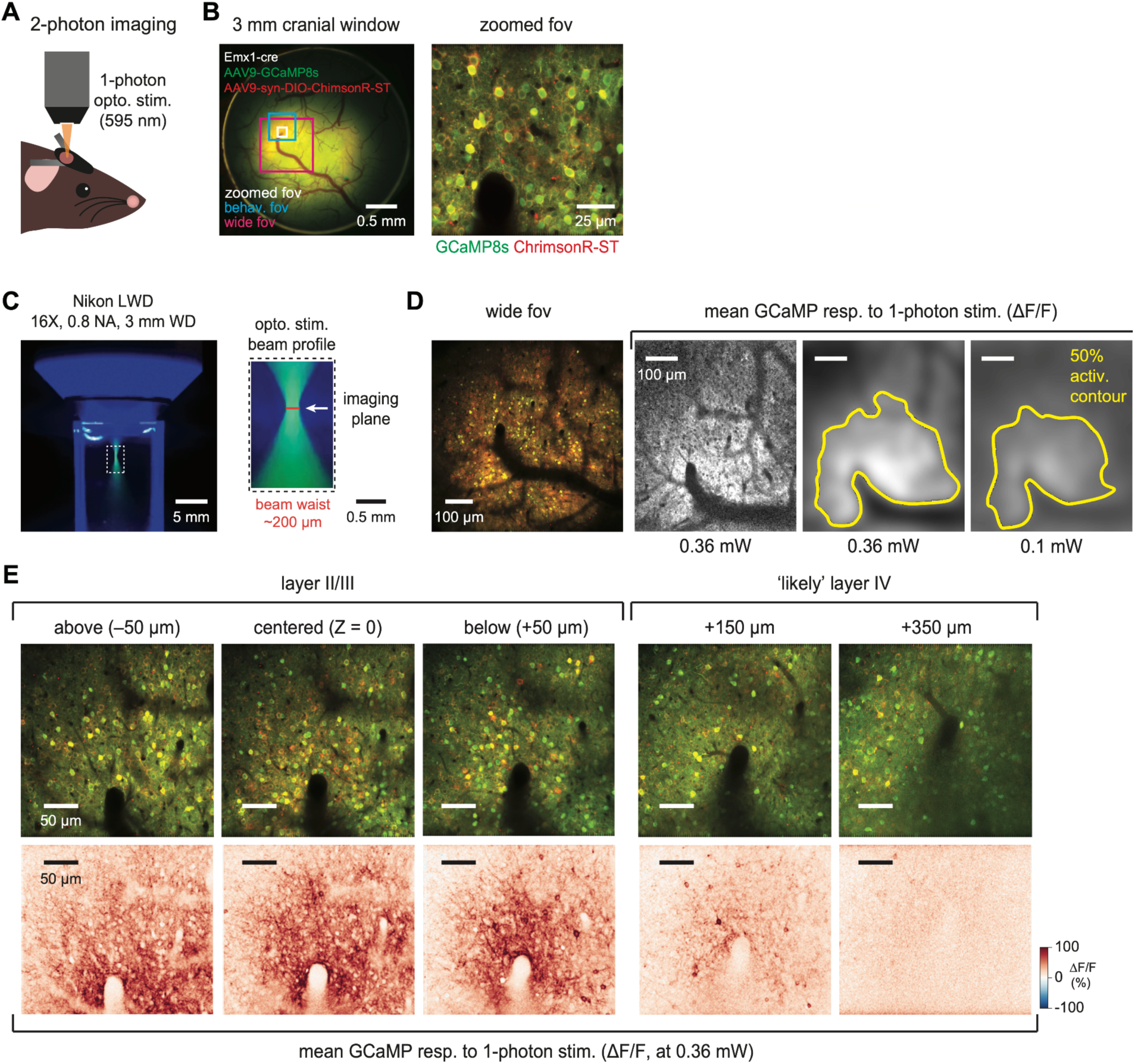
Spatial extent of 1-photon stimulation is confined in lateral and depth axes, Related to Figure 2. (**A**) Schematic of head-fixed 2-photon imaging, with 1-photon optogenetic stimulation delivered through the objective. (**B**) Widefield fluorescence through craniotomy. AAV9-GCaMP8s (green), stChrimsonR (red), and coexpression (yellow) in enlarged field-of-view (FOV). White, red, and blue boxes indicate FOVs for 2-photon imaging. Smallest FOV (white box) shown in the right panel, FOV for spatial measurement (red box) **D**, and FOV used for imaging (blue box) during behavior in **E**. (**C**) *In vitro* measurement of 1-photon stimulation beam profile in fluorescein solution. White dotted box shows the area zoomed on the right. White arrow shows the approximate imaging plane. Approximate beam waist is shown in red. (**D**) GCaMP and stChrimsonR expression in a wide 2-photon FOV. Right panels show mean ΔF/F response to 1-photon stimulation at two powers (0.36 and 0.1 mW). 50% activation contour is shown by the yellow outline. (**E**) Mean ΔF/F response (2-photon imaging with 1-photon optogenetic stimulation) at different depths in V1. Left panels show the responses of layer II/III neurons, center labeled panel (Z = 0, 150 µm below the cortical surface). Right panels show smaller neural responses in deeper cortical layers (+150 and +350 µm) labeled ‘likely layer IV’.

**Figure S6.**
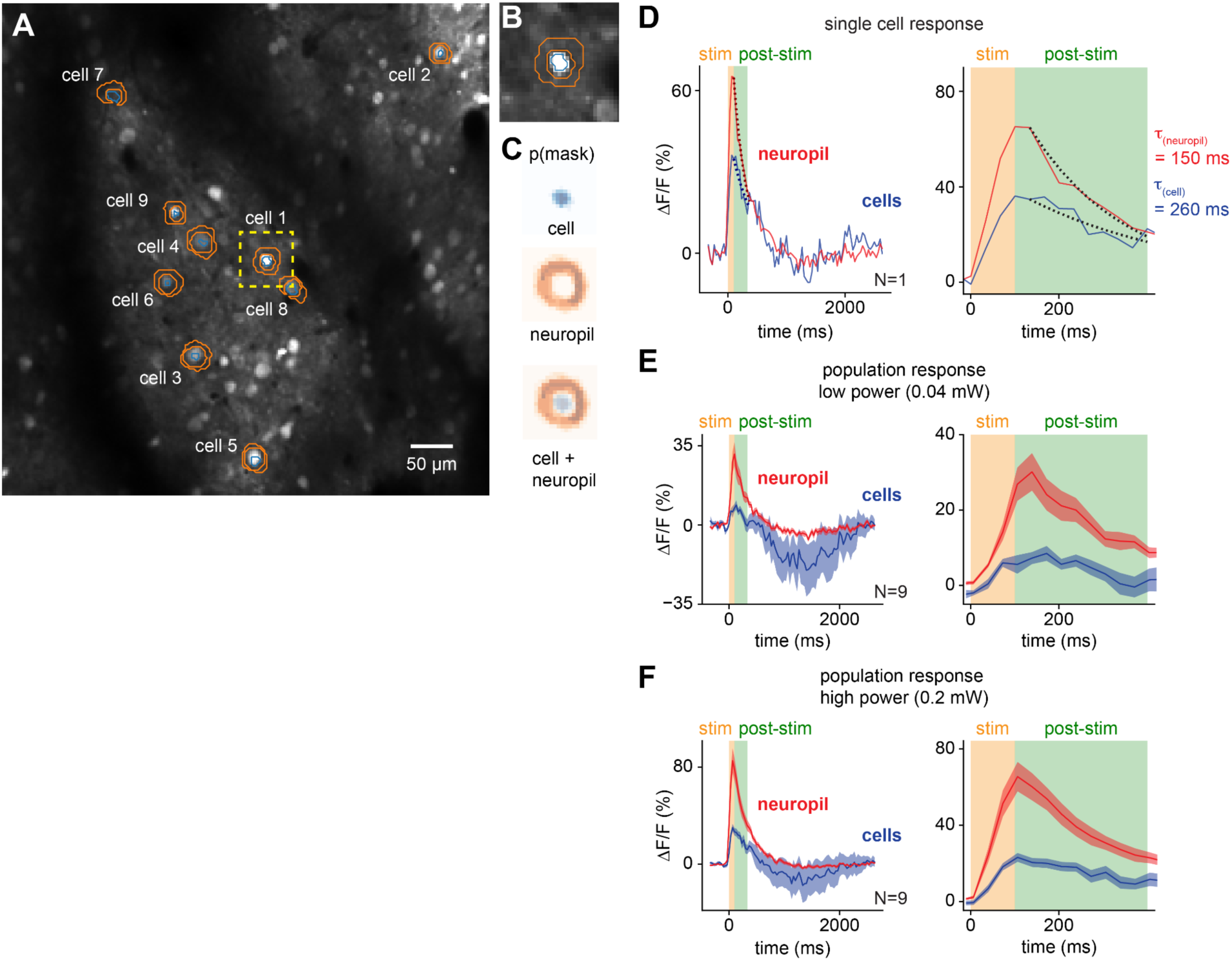
Cell soma and neuropil show distinct decay kinetics, Related to Figure 2. (**A**) Field-of-view image of 2-photon data collection from the first session in 1 animal. Nine relatively isolated cells were selected. An inner cell body mask was drawn (blue), and an annulus in the surrounding neuropil was drawn (orange), avoiding any nearby cell bodies. (**B**) Zoomed in view of cell 1, showing the cell body mask in blue and the neuropil annulus in orange. (**C**) Each mask was centered and averaged to produce mask probabilities for each compartment, cell in blue, and neuropil in orange. (**D**) Average response to 1-photon stimulation for an example cell for its surrounding neuropil (red) and the cell body (blue). Left: response zoomed in to the first 500 ms after stimulation onset. A single exponential decay was fit to each compartment and is depicted by the dotted lines, red and blue for neuropil and cell body respectively. Tau values represent the half decay times of the exponential decay fits. (**E**) Population average (N = 9) responses at 0.04 mW stimulation power, displayed analogously to **D**. (**F**) Same as **E** but for 0.2 mW stimulation power.

**Figure S7.**
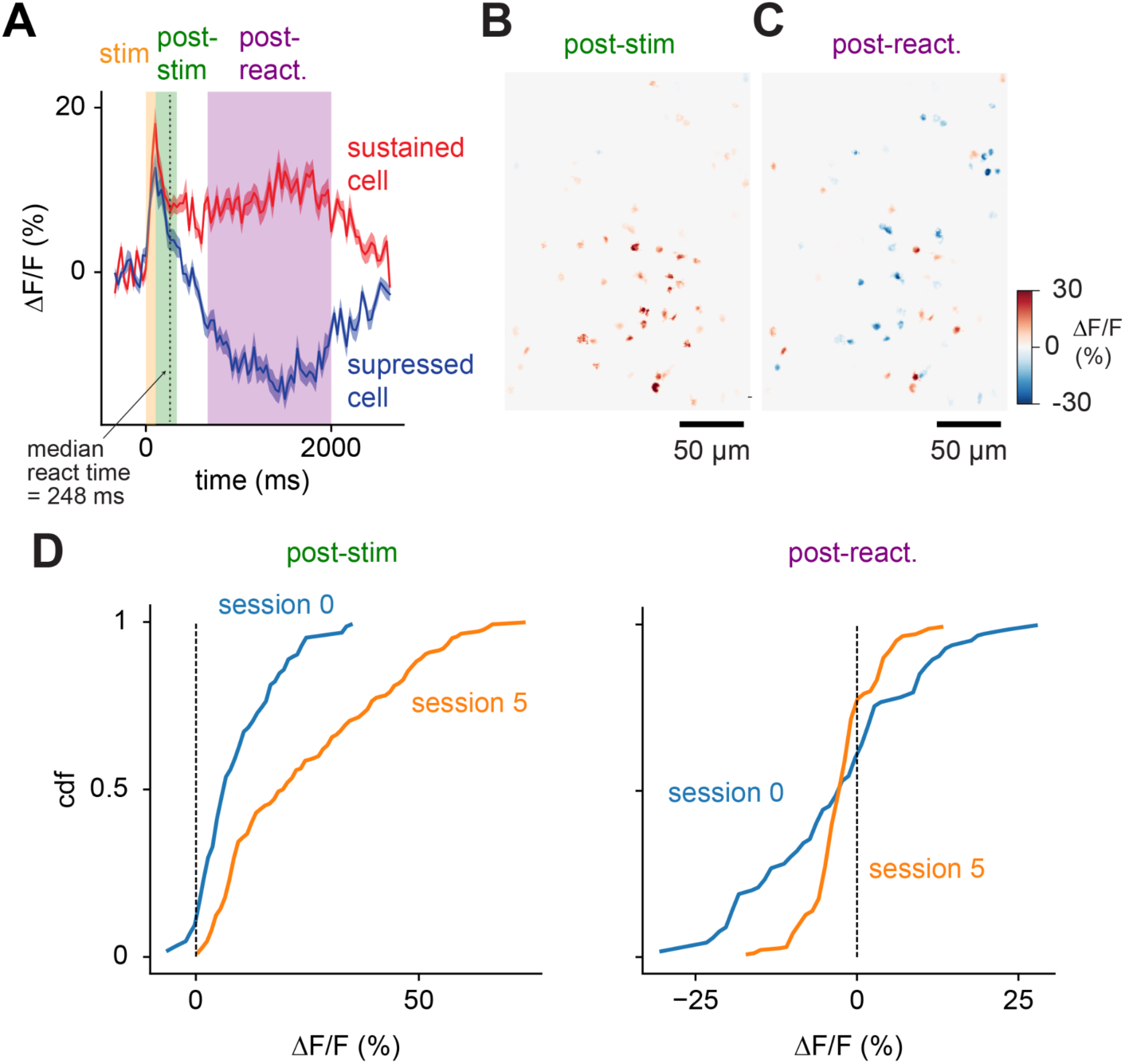
Behaviorally-relevant cell responses show elevated firing rates while post-decision responses show suppression, Related to Figure 2. (**A**) Population timecourses of response for cells that show a positive or negative response during the post-react period (sustained and suppressed, respectively). Analysis periods are highlighted: stimulation, orange; post-stimulation, green; and post-reaction, purple (see Methods). (**B**) Spatial distribution of average responses during the post-stim. period shows uniformly positive responses, while (**C**) distribution of responses during the post-react. period show salt-and-pepper sustained and suppressed responses. (**D**) Cumulative distributions of responses during the post-stim. period (left) and post-react. Period (right).

**Figure S8.**
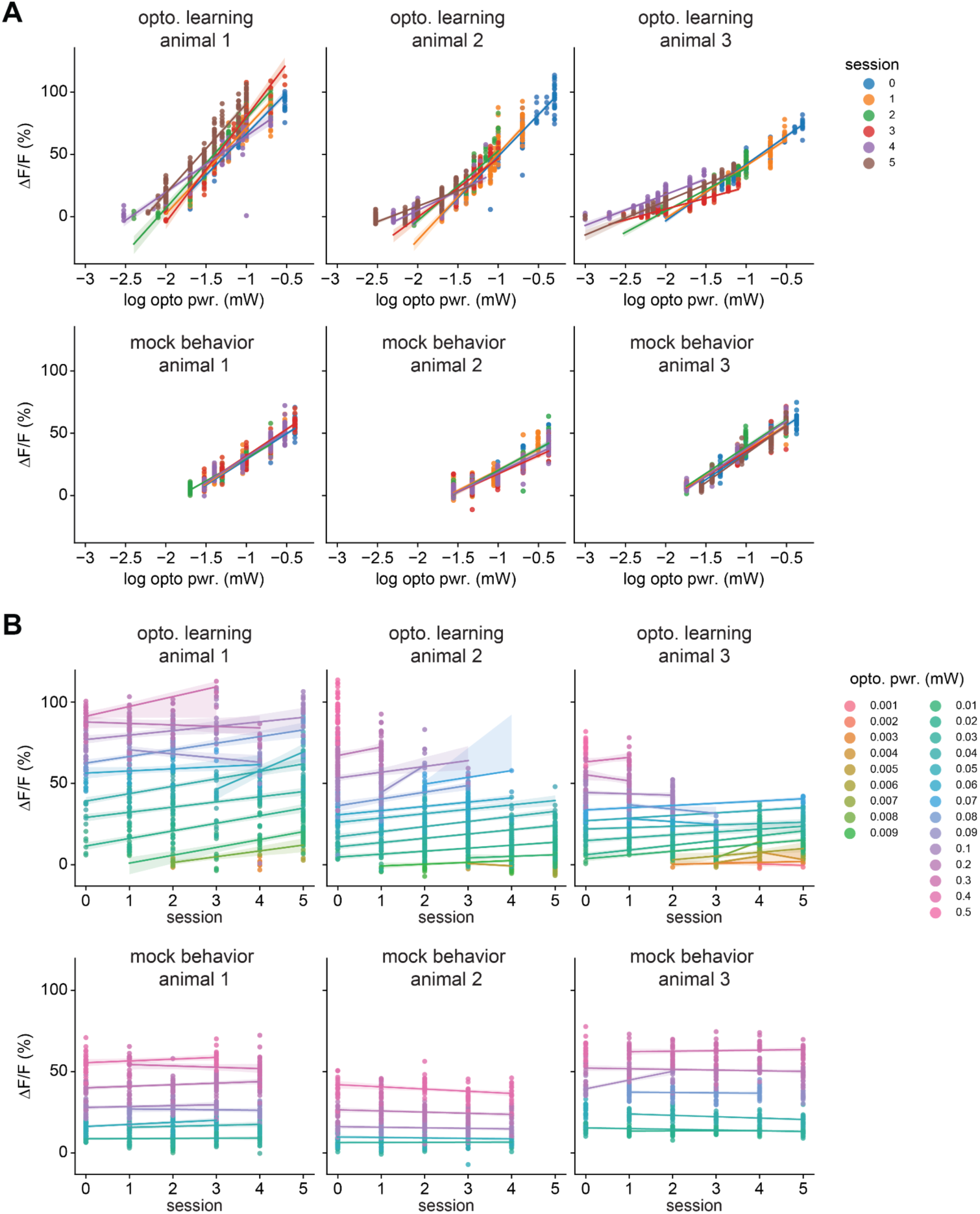
V1 amplification effect, all sessions, all animals, Related to Figure 2. (**A**) Linear regression model for testing for amplification effects between behavioral sessions (ROI-based population analysis shown in Figure 2C-J). OLS regression using all trials the stimulus was successfully detected (optogenetic learning animals: p = 1.73 x 10^-12^, N = 3 animals, 2633 trials; mock behavioral control animals: p = 0.26, N = 3, 1731 trials model: ΔF/F ∼ C(animal) * C(session) + stimulation_power_mw + trial_number + constant, where C(x) signifies a categorical or dummy variable). Treating power as a continuous variable did not change the results. In the three training animals, lines fit on each session (colors) moved leftward as learning progressed, signifying amplification. Within each session we found a small decrease in responses to stimulation (average ΔF/F change over 100 trials: −1.2% ΔF/F, 95% CI [-0.9 to 1.6]% ΔF/F, coeff. less than zero at p < 10^-13^, via linear regression over trials within day, estimated across animals and sessions, N = 3; Methods) (**B**) Comparison of amplification at each power across all behavioral sessions. Here, at many powers common across sessions (colors, lines), the slope of the corresponding line was positive, signifying amplification. We found a small decrease in responses to stimulation over the course of each experimental day.

**Figure S9.**
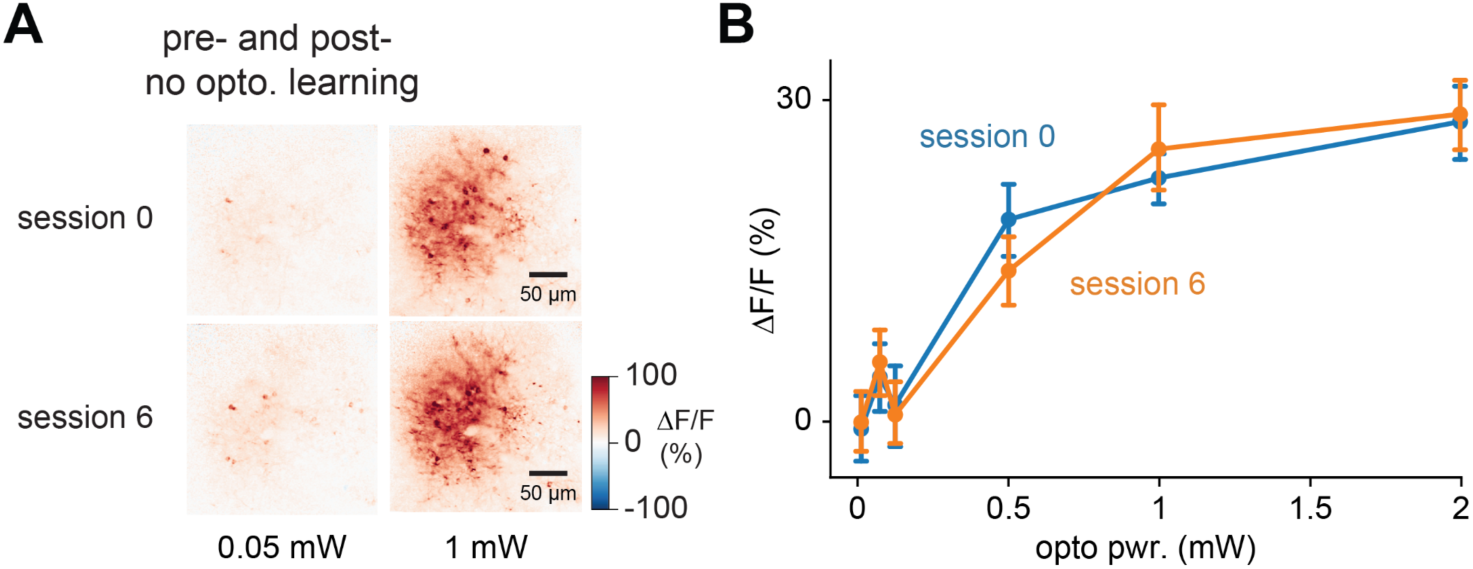
No amplification occurs for optogenetic stimulation delivered to V1 outside the learning context, Related to Figure 2. (**A**) Mean ΔF/F responses in an example animal to 0.05 and 1 mW of optogenetic stimulation delivered outside the behavioral context; animal was awake and alert but any motor responses were not reinforced (see Methods). 0.05 mW is near the average post-learning threshold power for optogenetic learning animals. 1 mW is a power level three times higher than the maximum used in training optogenetic learning animals. (**B**) No amplification occurs at any power level over seven sessions of optogenetic stimulation (example animal, session 0, blue, to session 6, orange, mean ± SEM). There was no significant change in response across N = 2 animals (session 0 vs. 6, via ANOVA/linear regression; t = 1.1, p = 0.27, also neither animal reaches significance alone, and treating power as a continuous or log-continuous variable did not change the results; see Methods for regression details).

**Figure S10.**
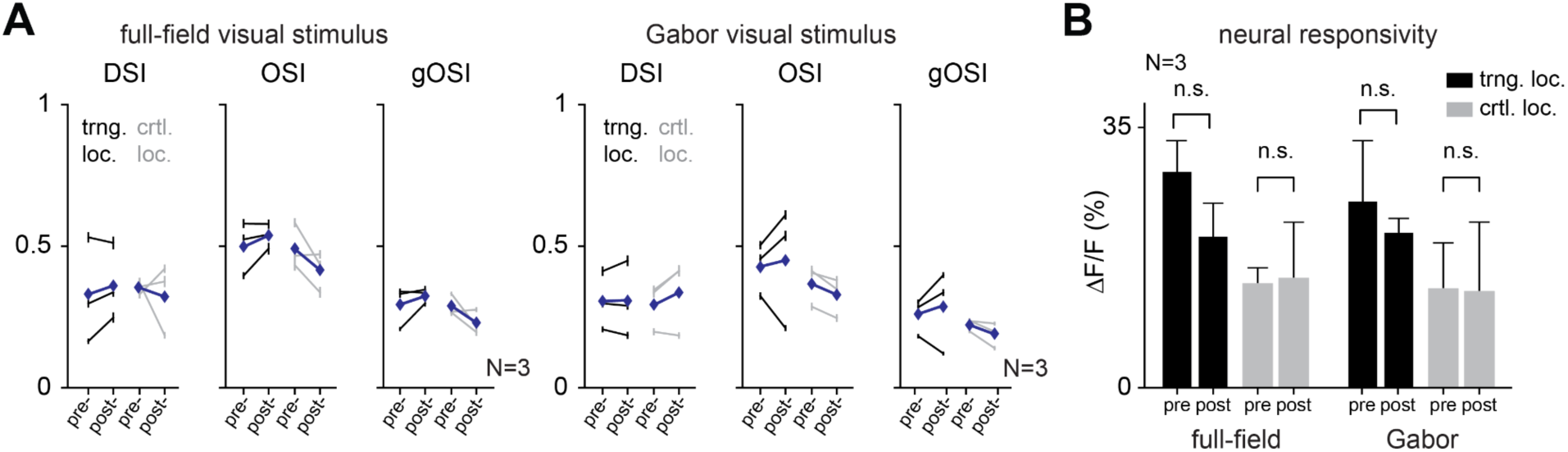
Visual response changes and neural responsivity before and after optogenetic learning, Related to Figure 3. (**A**) Visual response changes, pre- and post-learning, for individual animals. Cohort means shown by blue diamonds (mean ± SEM, N = 3). (**B**) Mean neural responsivity reveals no significant pre- versus post-learning changes at either the optogenetic training or control locations (mean ± SEM, N = 3).

**Figure S11.**
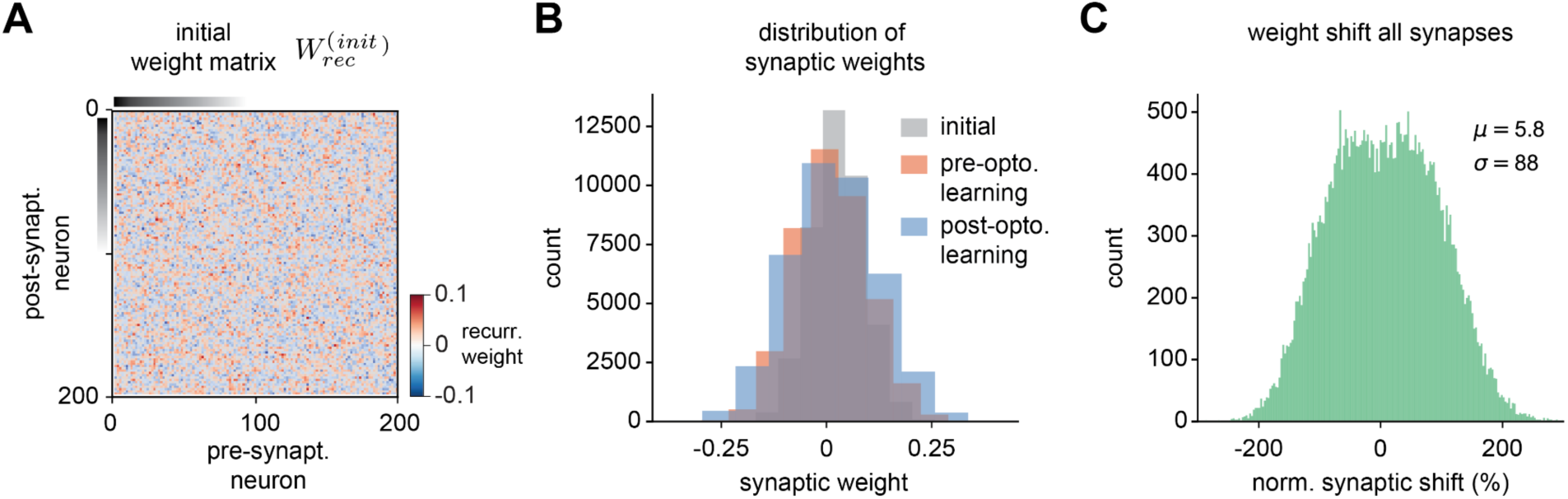
Synaptic weight matrix for initial configuration, and overall distributions of synaptic weights before and after training, Related to Figure 4. (**A**) Initial weight matrix before any training. Sampled from a random Gaussian distribution with mean zero (see Methods). (**B**) Distributions of synaptic weights, initial (gray), pre-opto. learning (red), and after training to amplify outputs (post-opto. learning, blue). (**C**) Distribution of synaptic weight changes (pre-opto. versus post-opto. learning), across all synapses, normalized to the mean pre-training weight.

## References

1. Chung, S., and Abbott, L.F. (2021). Neural population geometry: An approach for understanding biological and artificial neural networks. Curr. Opin. Neurobiol. 70, 137–144. 10.1016/j.conb.2021.10.010.

2. Gold, J.I., and Shadlen, M.N. (2007). The neural basis of decision making. Annu. Rev. Neurosci. 30, 535–574. 10.1146/annurev.neuro.29.051605.113038.

3. Wang, X.-J. (2012). Neural dynamics and circuit mechanisms of decision-making. Curr. Opin. Neurobiol. 22, 1039–1046. 10.1016/j.conb.2012.08.006.

4. Niell, C.M., and Scanziani, M. (2021). How Cortical Circuits Implement Cortical Computations: Mouse Visual Cortex as a Model. Annu. Rev. Neurosci. 44, 517–546. 10.1146/annurev-neuro-102320-085825.

5. Wu, Z., Litwin-Kumar, A., Shamash, P., Taylor, A., Axel, R., and Shadlen, M.N. (2020). Context-Dependent Decision Making in a Premotor Circuit. Neuron 106, 316–328.e6. 10.1016/j.neuron.2020.01.034.

6. Egger, S.W., and Lisberger, S.G. (2022). Neural structure of a sensory decoder for motor control. Nat. Commun. 13, 1829. 10.1038/s41467-022-29457-4.

7. Hengen, K.B., Lambo, M.E., Van Hooser, S.D., Katz, D.B., and Turrigiano, G.G. (2013). Firing rate homeostasis in visual cortex of freely behaving rodents. Neuron 80, 335–342. 10.1016/j.neuron.2013.08.038.

8. Malenka, R.C., and Bear, M.F. (2004). LTP and LTD: an embarrassment of riches. Neuron 44, 5–21. 10.1016/j.neuron.2004.09.012.

9. He, H.-Y., Hodos, W., and Quinlan, E.M. (2006). Visual deprivation reactivates rapid ocular dominance plasticity in adult visual cortex. J. Neurosci. 26, 2951–2955. 10.1523/JNEUROSCI.5554-05.2006.

10. Sawtell, N.B., Frenkel, M.Y., Philpot, B.D., Nakazawa, K., Tonegawa, S., and Bear, M.F. (2003). NMDA receptor-dependent ocular dominance plasticity in adult visual cortex. Neuron 38, 977–985. 10.1016/s0896-6273(03)00323-4.

11. Frégnac, Y., and Shulz, D.E. (1999). Activity-dependent regulation of receptive field properties of cat area 17 by supervised Hebbian learning. J. Neurobiol. 41, 69–82. 10.1002/(sici)1097-4695(199910)41:1[[[69::aid-neu10]]]3.0.co;2-1.

12. Schoups, A., Vogels, R., Qian, N., and Orban, G. (2001). Practising orientation identification improves orientation coding in V1 neurons. Nature 412, 549–553. 10.1038/35087601.

13. Schwartz, S., Maquet, P., and Frith, C. (2002). Neural correlates of perceptual learning: a functional MRI study of visual texture discrimination. Proc. Natl. Acad. Sci. U. S. A. 99, 17137–17142. 10.1073/pnas.242414599.

14. Li, W., Piëch, V., and Gilbert, C.D. (2008). Learning to link visual contours. Neuron 57, 442–451. 10.1016/j.neuron.2007.12.011.

15. Bao, M., Yang, L., Rios, C., He, B., and Engel, S.A. (2010). Perceptual learning increases the strength of the earliest signals in visual cortex. J. Neurosci. 30, 15080–15084. 10.1523/JNEUROSCI.5703-09.2010.

16. Ghose, G.M., Yang, T., and Maunsell, J.H.R. (2002). Physiological correlates of perceptual learning in monkey V1 and V2. J. Neurophysiol. 87, 1867–1888. 10.1152/jn.00690.2001.

17. Goltstein, P.M., Coffey, E.B.J., Roelfsema, P.R., and Pennartz, C.M.A. (2013). In vivo two-photon Ca2+ imaging reveals selective reward effects on stimulus-specific assemblies in mouse visual cortex. J. Neurosci. 33, 11540–11555. 10.1523/JNEUROSCI.1341-12.2013.

18. Poort, J., Khan, A.G., Pachitariu, M., Nemri, A., Orsolic, I., Krupic, J., Bauza, M., Sahani, M., Keller, G.B., Mrsic-Flogel, T.D., et al. (2015). Learning Enhances Sensory and Multiple Non-sensory Representations in Primary Visual Cortex. Neuron 86, 1478–1490. 10.1016/j.neuron.2015.05.037.

19. Goltstein, P.M., Reinert, S., Bonhoeffer, T., and Hübener, M. (2021). Mouse visual cortex areas represent perceptual and semantic features of learned visual categories. Nat. Neurosci. 24, 1441–1451. 10.1038/s41593-021-00914-5.

20. Yotsumoto, Y., Watanabe, T., and Sasaki, Y. (2008). Different dynamics of performance and brain activation in the time course of perceptual learning. Neuron 57, 827–833. 10.1016/j.neuron.2008.02.034.

21. Jurjut, O., Georgieva, P., Busse, L., and Katzner, S. (2017). Learning Enhances Sensory Processing in Mouse V1 before Improving Behavior. J. Neurosci. 37, 6460–6474. 10.1523/JNEUROSCI.3485-16.2017.

22. Marshel, J.H., Kim, Y.S., Machado, T.A., Quirin, S., Benson, B., Kadmon, J., Raja, C., Chibukhchyan, A., Ramakrishnan, C., Inoue, M., et al. (2019). Cortical layer-specific critical dynamics triggering perception. Science 365. 10.1126/science.aaw5202.

23. Henschke, J.U., Dylda, E., Katsanevaki, D., Dupuy, N., Currie, S.P., Amvrosiadis, T., Pakan, J.M.P., and Rochefort, N.L. (2020). Reward Association Enhances Stimulus-Specific Representations in Primary Visual Cortex. Curr. Biol. 30, 1866–1880.e5. 10.1016/j.cub.2020.03.018.

24. Khan, A.G., Poort, J., Chadwick, A., Blot, A., Sahani, M., Mrsic-Flogel, T.D., and Hofer, S.B. (2018). Distinct learning-induced changes in stimulus selectivity and interactions of GABAergic interneuron classes in visual cortex. Nat. Neurosci. 21, 851–859. 10.1038/s41593-018-0143-z.

25. Boynton, G.M., and Finney, E.M. (2003). Orientation-Specific Adaptation in Human Visual Cortex. J. Neurosci. 23, 8781–8787. 10.1523/jneurosci.23-25-08781.2003.

26. Yang, T., and Maunsell, J.H.R. (2004). The effect of perceptual learning on neuronal responses in monkey visual area V4. J. Neurosci. 24, 1617–1626. 10.1523/JNEUROSCI.4442-03.2004.

27. Dosher, B., and Lu, Z.-L. (2017). Visual Perceptual Learning and Models. Annu. Rev. Vis. Sci. 3, 343–363. 10.1146/annurev-vision-102016-061249.

28. Schmolesky, M.T., Wang, Y., Hanes, D.P., Thompson, K.G., Leutgeb, S., Schall, J.D., and Leventhal, A.G. (1998). Signal timing across the macaque visual system. J. Neurophysiol. 79, 3272–3278. 10.1152/jn.1998.79.6.3272.

29. Steinmetz, N.A., Zatka-Haas, P., Carandini, M., and Harris, K.D. (2019). Distributed coding of choice, action and engagement across the mouse brain. Nature 576, 266–273. 10.1038/s41586-019-1787-x.

30. Zatka-Haas, P., Steinmetz, N.A., Carandini, M., and Harris, K.D. (2021). Sensory coding and the causal impact of mouse cortex in a visual decision. eLife 10. 10.7554/eLife.63163.

31. Liang, L., Fratzl, A., Reggiani, J.D.S., El Mansour, O., Chen, C., and Andermann, M.L. (2020). Retinal Inputs to the Thalamus Are Selectively Gated by Arousal. Curr. Biol. 30, 3923–3934.e9. 10.1016/j.cub.2020.07.065.

32. Jazayeri, M., and Afraz, A. (2017). Navigating the Neural Space in Search of the Neural Code. Neuron 93, 1003–1014. 10.1016/j.neuron.2017.02.019.

33. Sadtler, P.T., Quick, K.M., Golub, M.D., Chase, S.M., Ryu, S.I., Tyler-Kabara, E.C., Yu, B.M., and Batista, A.P. (2014). Neural constraints on learning. Nature 512, 423–426. 10.1038/nature13665.

34. Murphy, B.K., and Miller, K.D. (2009). Balanced amplification: a new mechanism of selective amplification of neural activity patterns. Neuron 61, 635–648. 10.1016/j.neuron.2009.02.005.

35. Goldman, M.S. (2009). Memory without feedback in a neural network. Neuron 61, 621–634. 10.1016/j.neuron.2008.12.012.

36. Hennequin, G., Vogels, T.P., and Gerstner, W. (2012). Non-normal amplification in random balanced neuronal networks. Phys. Rev. E Stat. Nonlin. Soft Matter Phys. 86, 011909. 10.1103/PhysRevE.86.011909.

37. Pégard, N.C., Mardinly, A.R., Oldenburg, I.A., Sridharan, S., Waller, L., and Adesnik, H. (2017). Three-dimensional scanless holographic optogenetics with temporal focusing (3D-SHOT). Nature Communications 8. 10.1038/s41467-017-01031-3.

38. Cardin, J.A., Carlén, M., Meletis, K., Knoblich, U., Zhang, F., Deisseroth, K., Tsai, L.-H., and Moore, C.I. (2010). Targeted optogenetic stimulation and recording of neurons in vivo using cell-type-specific expression of Channelrhodopsin-2. Nat. Protocols 5, 247–254. 10.1038/nprot.2009.228.

39. Sohal, D.S., Nghiem, M., Crackower, M.A., Witt, S.A., Kimball, T.R., Tymitz, K.M., Penninger, J.M., and Molkentin, J.D. (2001). Temporally regulated and tissue-specific gene manipulations in the adult and embryonic heart using a tamoxifen-inducible Cre protein. Circ. Res. 89, 20–25. 10.1161/hh1301.092687.

40. Dana, H., Sun, Y., Mohar, B., Hulse, B.K., Kerlin, A.M., Hasseman, J.P., Tsegaye, G., Tsang, A., Wong, A., Patel, R., et al. (2019). High-performance calcium sensors for imaging activity in neuronal populations and microcompartments. Nat. Methods 16, 649–657. 10.1038/s41592-019-0435-6.

41. Kügler, S., Kilic, E., and Bähr, M. (2003). Human synapsin 1 gene promoter confers highly neuron-specific long-term transgene expression from an adenoviral vector in the adult rat brain depending on the transduced area. Gene Therapy 10, 337–347. 10.1038/sj.gt.3301905.

42. Zhang, Y., Rózsa, M., Bushey, D., Zheng, J., Reep, D., Broussard, G.J., Tsang, A., Tsegaye, G., Patel, R., Narayan, S., et al. (2020). jGCaMP8 fast genetically encoded calcium indicators. Janelia Research Campus 10, 13148243.

43. Histed, M.H., and Maunsell, J.H.R. (2014). Cortical neural populations can guide behavior by integrating inputs linearly, independent of synchrony. Proc. Natl. Acad. Sci. U. S. A. 111, E178–E187. 10.1073/pnas.1318750111.

44. Macmillan, N.A., and Douglas Creelman, C. (2005). Detection Theory: A User’s Guide (Lawrence Erlbaum Associates).

45. Dalgleish, H.W., Russell, L.E., Packer, A.M., Roth, A., Gauld, O.M., Greenstreet, F., Thompson, E.J., and Häusser, M. (2020). How many neurons are sufficient for perception of cortical activity? eLife 9. 10.7554/eLife.58889.

46. O’Rawe, J.F., Zhou, Z., Li, A.J., LaFosse, P.K., Goldbach, H.C., and Histed, M.H. (2022). Excitation creates a distributed pattern of cortical suppression due to varied recurrent input. bioRxiv, 2022.08.31.505844. 10.1101/2022.08.31.505844.

47. Brunel, N. (2000). Dynamics of sparsely connected networks of excitatory and inhibitory spiking neurons. J. Comput. Neurosci. 8, 183–208. 10.1023/a:1008925309027.

48. Sanzeni, A., Histed, M.H., and Brunel, N. (2022). Emergence of Irregular Activity in Networks of Strongly Coupled Conductance-Based Neurons. Phys. Rev. X 12, 011044. 10.1103/PhysRevX.12.011044.

49. Ahmadian, Y., and Miller, K.D. (2021). What is the dynamical regime of cerebral cortex? Neuron 109, 3373–3391. 10.1016/j.neuron.2021.07.031.

50. Sanzeni, A., Akitake, B., Goldbach, H.C., Leedy, C.E., Brunel, N., and Histed, M.H. (2020). Inhibition stabilization is a widespread property of cortical networks. eLife 9. 10.7554/elife.54875.

51. Destexhe, A., and Paré, D. (1999). Impact of network activity on the integrative properties of neocortical pyramidal neurons in vivo. J. Neurophysiol. 81, 1531–1547. 10.1152/jn.1999.81.4.1531.

52. Mainen, Z.F., and Sejnowski, T.J. (1995). Reliability of spike timing in neocortical neurons. Science 268, 1503–1506. 10.1126/science.7770778.

53. Friedrich, J., Zhou, P., and Paninski, L. (2017). Fast online deconvolution of calcium imaging data. PLoS Comput. Biol. 13, e1005423. 10.1371/journal.pcbi.1005423.

54. Stern, M., Shea-Brown, E., and Witten, D. (2020). Inferring the Spiking Rate of a Population of Neurons from Wide-Field Calcium Imaging. bioRxiv, 2020.02.01.930040. 10.1101/2020.02.01.930040.

55. Pachitariu, M., Stringer, C., Dipoppa, M., Schröder, S., Federico Rossi, L., Dalgleish, H., Carandini, M., and Harris, K.D. (2017). Suite2p: beyond 10,000 neurons with standard two-photon microscopy. bioRxiv, 061507. 10.1101/061507.

56. Hubel, D.H., and Wiesel, T.N. (1962). Receptive fields, binocular interaction and functional architecture in the cat’s visual cortex. J. Physiol. 160, 106–154. 10.1113/jphysiol.1962.sp006837.

57. Cano, M., Bezdudnaya, T., Swadlow, H.A., and Alonso, J.-M. (2006). Brain state and contrast sensitivity in the awake visual thalamus. Nat. Neurosci. 9, 1240–1242. 10.1038/nn1760.

58. Sadagopan, S., and Ferster, D. (2012). Feedforward origins of response variability underlying contrast invariant orientation tuning in cat visual cortex. Neuron 74, 911–923. 10.1016/j.neuron.2012.05.007.

59. Kelly, S.T., Kremkow, J., Jin, J., Wang, Y., Wang, Q., Alonso, J.-M., and Stanley, G.B. (2014). The role of thalamic population synchrony in the emergence of cortical feature selectivity. PLoS Comput. Biol. 10, e1003418. 10.1371/journal.pcbi.1003418.

60. Deitch, D., Rubin, A., and Ziv, Y. (2021). Representational drift in the mouse visual cortex. Curr. Biol. 31, 4327–4339.e6. 10.1016/j.cub.2021.07.062.

61. Marks, T.D., and Goard, M.J. (2021). Stimulus-dependent representational drift in primary visual cortex. Nat. Commun. 12, 5169. 10.1038/s41467-021-25436-3.

62. Moreno-Bote, R., Beck, J., Kanitscheider, I., Pitkow, X., Latham, P., and Pouget, A. (2014). Information-limiting correlations. Nat. Neurosci. 17, 1410–1417. 10.1038/nn.3807.

63. Pancholi, R., Ryan, L., and Peron, S. (2021). Sensory cortical dynamics during optical microstimulation training. bioRxiv, 2021.12.17.473191. 10.1101/2021.12.17.473191.

64. Gill, J.V., Lerman, G.M., Zhao, H., Stetler, B.J., Rinberg, D., and Shoham, S. (2020). Precise Holographic Manipulation of Olfactory Circuits Reveals Coding Features Determining Perceptual Detection. Neuron 108, 382–393.e5. 10.1016/j.neuron.2020.07.034.

65. Histed, M.H., Bonin, V., and Reid, R.C. (2009). Direct activation of sparse, distributed populations of cortical neurons by electrical microstimulation. Neuron 63, 508–522. 10.1016/j.neuron.2009.07.016.

66. Histed, M.H., Ni, A.M., and Maunsell, J.H.R. (2013). Insights into cortical mechanisms of behavior from microstimulation experiments. Prog. Neurobiol. 103, 115–130. 10.1016/j.pneurobio.2012.01.006.

67. Ni, A.M., and Maunsell, J.H.R. (2010). Microstimulation reveals limits in detecting different signals from a local cortical region. Curr. Biol. 20, 824–828. 10.1016/j.cub.2010.02.065.

68. Doty, R.W. (1969). Electrical stimulation of the brain in behavioral context. Annu. Rev. Psychol. 20, 289–320. 10.1146/annurev.ps.20.020169.001445.

69. Doron, G., and Brecht, M. (2015). What single-cell stimulation has told us about neural coding. Philos. Trans. R. Soc. Lond. B Biol. Sci. 370, 20140204. 10.1098/rstb.2014.0204.

70. Luis-Islas, J., Luna, M., Floran, B., and Gutierrez, R. (2022). Optoception: Perception of Optogenetic Brain Perturbations. eNeuro 9. 10.1523/ENEURO.0216-22.2022.

71. Kohn, A., Coen-Cagli, R., Kanitscheider, I., and Pouget, A. (2016). Correlations and Neuronal Population Information. Annu. Rev. Neurosci. 10.1146/annurev-neuro-070815-013851.

72. Brashers-Krug, T., Shadmehr, R., and Bizzi, E. (1996). Consolidation in human motor memory. Nature 382, 252–255. 10.1038/382252a0.

73. Krakauer, J.W., and Shadmehr, R. (2006). Consolidation of motor memory. Trends Neurosci. 29, 58–64. 10.1016/j.tins.2005.10.003.

74. Pons, T.P., Garraghty, P.E., Ommaya, A.K., Kaas, J.H., Taub, E., and Mishkin, M. (1991). Massive cortical reorganization after sensory deafferentation in adult macaques. Science 252, 1857–1860. 10.1126/science.1843843.

75. Gilbert, C.D., and Li, W. (2012). Adult visual cortical plasticity. Neuron 75, 250–264. 10.1016/j.neuron.2012.06.030.

76. Alejandre-García, T., Kim, S., Pérez-Ortega, J., and Yuste, R. (2022). Intrinsic excitability mechanisms of neuronal ensemble formation. Elife 11. 10.7554/eLife.77470.

77. Sadeh, S., and Clopath, C. (2020). Theory of neuronal perturbome in cortical networks. Proc. Natl. Acad. Sci. U. S. A. 117, 26966–26976. 10.1073/pnas.2004568117.

78. Carrillo-Reid, L., and Yuste, R. (2020). Playing the piano with the cortex: role of neuronal ensembles and pattern completion in perception and behavior. Curr. Opin. Neurobiol. 64, 89–95. 10.1016/j.conb.2020.03.014.

79. Peters, A.J., Liu, H., and Komiyama, T. (2017). Learning in the Rodent Motor Cortex. Annu. Rev. Neurosci. 40, 77–97. 10.1146/annurev-neuro-072116-031407.

80. Hedrick, N.G., Lu, Z., Bushong, E., Singhi, S., Nguyen, P., Magaña, Y., Jilani, S., Lim, B.K., Ellisman, M., and Komiyama, T. (2022). Learning binds new inputs into functional synaptic clusters via spinogenesis. Nature Neuroscience 25, 726–737. 10.1038/s41593-022-01086-6.

81. Liu, B., Seay, M.J., and Buonomano, D.V. (2023). Creation of Neuronal Ensembles and Cell-Specific Homeostatic Plasticity through Chronic Sparse Optogenetic Stimulation. J. Neurosci. 43, 82–92. 10.1523/JNEUROSCI.1104-22.2022.

82. Keck, T., Keller, G.B., Jacobsen, R.I., Eysel, U.T., Bonhoeffer, T., and Hübener, M. (2013). Synaptic scaling and homeostatic plasticity in the mouse visual cortex in vivo. Neuron 80, 327–334. 10.1016/j.neuron.2013.08.018.

83. Swanson, O.K., and Maffei, A. (2019). From Hiring to Firing: Activation of Inhibitory Neurons and Their Recruitment in Behavior. Front. Mol. Neurosci. 12, 168. 10.3389/fnmol.2019.00168.

84. Trachtenberg, J.T. (2015). Competition, inhibition, and critical periods of cortical plasticity. Curr. Opin. Neurobiol. 35, 44–48. 10.1016/j.conb.2015.06.006.

85. Heimel, J.A., van Versendaal, D., and Levelt, C.N. (2011). The role of GABAergic inhibition in ocular dominance plasticity. Neural Plast. 2011, 391763. 10.1155/2011/391763.

86. Fagiolini, M., and Hensch, T.K. (2000). Inhibitory threshold for critical-period activation in primary visual cortex. Nature 404, 183–186. 10.1038/35004582.

87. Fagiolini, M., Fritschy, J.-M., Löw, K., Möhler, H., Rudolph, U., and Hensch, T.K. (2004). Specific GABAA circuits for visual cortical plasticity. Science 303, 1681–1683. 10.1126/science.1091032.

88. Carcea, I., and Froemke, R.C. (2013). Chapter 3 - Cortical Plasticity, Excitatory–Inhibitory Balance, and Sensory Perception. In Progress in Brain Research, M. M. Merzenich, M. Nahum, and T. M. Van Vleet, eds. (Elsevier), pp. 65–90. 10.1016/B978-0-444-63327-9.00003-5.

89. Reichelt, A.C., Hare, D.J., Bussey, T.J., and Saksida, L.M. (2019). Perineuronal Nets: Plasticity, Protection, and Therapeutic Potential. Trends Neurosci. 42, 458–470. 10.1016/j.tins.2019.04.003.

90. Hylin, M.J., Orsi, S.A., Moore, A.N., and Dash, P.K. (2013). Disruption of the perineuronal net in the hippocampus or medial prefrontal cortex impairs fear conditioning. Learn. Mem. 20, 267–273. 10.1101/lm.030197.112.

91. Banerjee, S.B., Gutzeit, V.A., Baman, J., Aoued, H.S., Doshi, N.K., Liu, R.C., and Ressler, K.J. (2017). Perineuronal Nets in the Adult Sensory Cortex Are Necessary for Fear Learning. Neuron 95, 169–179.e3. 10.1016/j.neuron.2017.06.007.

92. Le Naour, F., Hohenkirk, L., Grolleau, A., Misek, D.E., Lescure, P., Geiger, J.D., Hanash, S., and Beretta, L. (2001). Profiling changes in gene expression during differentiation and maturation of monocyte-derived dendritic cells using both oligonucleotide microarrays and proteomics. J. Biol. Chem. 276, 17920–17931. 10.1074/jbc.M100156200.

93. Sorg, B.A., Berretta, S., Blacktop, J.M., Fawcett, J.W., Kitagawa, H., Kwok, J.C.F., and Miquel, M. (2016). Casting a Wide Net: Role of Perineuronal Nets in Neural Plasticity. J. Neurosci. 36, 11459–11468. 10.1523/JNEUROSCI.2351-16.2016.

94. Balmer, T.S., Carels, V.M., Frisch, J.L., and Nick, T.A. (2009). Modulation of perineuronal nets and parvalbumin with developmental song learning. J. Neurosci. 29, 12878–12885. 10.1523/JNEUROSCI.2974-09.2009.

95. Gu, Y., Tran, T., Murase, S., Borrell, A., Kirkwood, A., and Quinlan, E.M. (2016). Neuregulin-dependent regulation of fast-spiking interneuron excitability controls the timing of the critical period. J. Neurosci. 36, 10285–10295. 10.1523/JNEUROSCI.4242-15.2016.

96. Hensch, T.K. (2005). Critical period plasticity in local cortical circuits. Nat. Rev. Neurosci. 6, 877–888. 10.1038/nrn1787.

97. Katz, L.C., and Crowley, J.C. (2002). Development of cortical circuits: lessons from ocular dominance columns. Nat. Rev. Neurosci. 3, 34–42. 10.1038/nrn703.

98. Desai, N.S., Cudmore, R.H., Nelson, S.B., and Turrigiano, G.G. (2002). Critical periods for experience-dependent synaptic scaling in visual cortex. Nat. Neurosci. 5, 783–789. 10.1038/nn878.

99. Gorski, J.A., Talley, T., Qiu, M., Puelles, L., Rubenstein, J.L.R., and Jones, K.R. (2002). Cortical excitatory neurons and glia, but not GABAergic neurons, are produced in the Emx1-expressing lineage. J. Neurosci. 22, 6309–6314. 20026564.

100. Goldbach, H.C., Akitake, B., Leedy, C.E., and Histed, M.H. (2021). Performance in even a simple perceptual task depends on mouse secondary visual areas. eLife 10. 10.7554/eLife.62156.

101. Histed, M.H., Carvalho, L.A., and Maunsell, J.H.R. (2012). Psychophysical measurement of contrast sensitivity in the behaving mouse. J. Neurophysiol. 107, 758–765. 10.1152/jn.00609.2011.

102. Macmillan, N.A., and Creelman, C.D. (2004). Detection theory: A user’s guide (Psychology press).

103. Kerlin, A., Mohar, B., Flickinger, D., MacLennan, B.J., Dean, M.B., Davis, C., Spruston, N., and Svoboda, K. (2019). Functional clustering of dendritic activity during decision-making. eLife 8. 10.7554/eLife.46966.

104. Giovannucci, A., Friedrich, J., Gunn, P., Kalfon, J., Brown, B.L., Koay, S.A., Taxidis, J., Najafi, F., Gauthier, J.L., Zhou, P., et al. (2019). CaImAn an open source tool for scalable calcium imaging data analysis. eLife 8. 10.7554/eLife.38173.

105. Swindale, N.V. (1998). Orientation tuning curves: empirical description and estimation of parameters. Biol. Cybern. 78, 45–56. 10.1007/s004220050411.

106. Kondo, S., and Ohki, K. (2016). Laminar differences in the orientation selectivity of geniculate afferents in mouse primary visual cortex. Nat. Neurosci. 19, 316–319. 10.1038/nn.4215.

107. Sompolinsky, H., Crisanti, A., and Sommers, H.J. (1988). Chaos in random neural networks. Phys. Rev. Lett. 61, 259–262. 10.1103/PhysRevLett.61.259.

108. Kingma, D.P., and Ba, J. (2014). Adam: A Method for Stochastic Optimization. arXiv [cs.LG].

109. Paszke, A., Gross, S., Massa, F., Lerer, A., Bradbury, J., Chanan, G., Killeen, T., Lin, Z., Gimelshein, N., Antiga, L., et al. (2019). PyTorch: An imperative style, high-performance deep learning library. arXiv [cs.LG].

